# ASigNTF: Agnostic Signature using NTF – a universal agnostic strategy to estimate cell-types abundance from transcriptomic datasets

**DOI:** 10.1101/2021.02.04.429589

**Authors:** Paul Fogel, Galina Boldina, Corinne Rocher, Charles Bettembourg, George Luta, Franck Augé

## Abstract

**Background:** Molecular signatures for deconvolution of immune cell types have been proposed, based on a methodology that relies on the biological classification of the cell types being studied. When working with less known biological material, a data-driven approach is needed to uncover the underlying classes and construct ad hoc signatures.

**Results:** We introduce a new approach, ASigNTF: Agnostic Signature using Non-negative Tensor Factorization, to perform the deconvolution of cell types from transcriptomics data (RNAseq and microarray). ASigNTF, which is based on two complementary statistical/mathematical tools: non-negative tensor factorization (for dimensionality reduction) and the Herfindahl-Hirschman index (for signature selection), can be applied to any type of tissue as long as transcriptomic data on isolated cells is available. As a direct result of the new method, we propose two new signatures for the deconvolution of immune cell types, one consisting of a relatively small set of 415 genes, which is more compatible with microarray platforms, and a larger set of 915 genes. Using external datasets, our two signatures outperform the CIBERSORT LM22 signature in deconvolution of RNA-seq data. Our signature with 415 genes allows to recognize a larger number of cell types compared to the ABIS microarray signature.

**Conclusions:** The paper proposes a new method, ASigNTF; applies the method, and also provides a software implementation that allows to identify molecular signatures for deconvolution of complex tissues and specifically up to 16 immune cell types from micro-array or RNA-seq data.

**Highlights:**

- Several signatures of immune cell types have been proposed, which follow a methodology deeply rooted in the known biological classification of the investigated cell types.
- When working with less known biological material, a more agnostic, data-driven approach is required to uncover the underlying classes and construct ad hoc signatures.
- We present ASigNTF, a new agnostic approach to cell type classification and signature selection supported by an application software.
- We discuss the results of benchmarking our proposed signatures, ABIS-seq and CIBERSORT on external datasets.

## Introduction

The identification of cell type components and their respective proportions from a heterogeneous biological sample can be achieved numerically by a so-called *deconvolution* process. This *in-silico* enumeration is fast and does not require complex biological experiments (e.g. flow cytometry, FACS). This is an important field of research as it can help to gain insights into the underlying biology without additional experiments or the need to perform a new analysis on existing datasets.

Deconvolution methods only work for cell types that have a detectable and specific signal pattern, which is typically delivered by subgroups of *marker genes* that constitute the cell types *signature*. Consequently, cell types that are close to each other should be grouped together. This step is often based on previous biological knowledge and can be complicated for complex cell types such as Peripheral Blood Mononuclear Cells, PBMC, as several cell type combinations are possible.

Along with an optimal grouping of cell types, highly detectable and specific marker genes are the key to the good performance of deconvolution methods. A traditional approach to identifying marker genes is the differential analysis between each cell type and the pool of all other cell types [1], or the second or third highest expressed group [2]. Both approaches have a number of limitations, just to name a few: Depending on the method used to perform the differential analysis, low-expressed genes may be penalized so they do not appear to be the most significant [3] [4]; the list of differentially expressed genes for each cell type needs to be further filtered by using an arbitrary cut-off for fold changes; additional filtering techniques – such as minimizing a condition number [5] – are required to further remove genes with low specificity [1] [2] [6]. The adjustment of different parameters at each step of the analysis workflow makes the identification of marker genes tedious and time-consuming.

Non-Negative Matrix Factorization, NMF [7] [8], or closely related techniques have been successfully used for blind identification of patterns of gene expression – molecular signatures that characterize different biological components [9] [10] [11] [12]. Specifically, bulk samples, mixtures of different populations, are analyzed to jointly estimate the gene expression of pure populations and the mixing proportions.

Here we introduce the Non-Negative Tensor Factorization, NTF [13], which extends the NMF to the analysis of more complex data. In this study, NTF, as opposed to blind identification methods, is applied to purified flow cytometric samples corresponding to known cell types, using the ABIS data set published by Monaco et al [1]. NTF allows the grouping of closely related cell types without previous knowledge of cell biology to make them suitable for deconvolution. Since we have access to the purified samples, we propose two simple rules for identifying highly specific marker genes, two cell type signatures and a simple deconvolution algorithm using non-negative least squares linear regression [14].

We evaluate our results using several public datasets focused on PBMC and whole blood tissues and compare them with those obtained by available signatures and software, namely ABIS-seq, ABIS-microarray and CIBERSORT [1] [15].

## Results

To address the limitations of the currently existing methods described in the introduction, we are proposing a new methodology, ASigNTF: Agnostic Signature using NTF. A summary description of the new method can be found in Box 1.

### Box 1. Summary description of our new methodology to construct highly specific cell type signatures and to perform deconvolution.

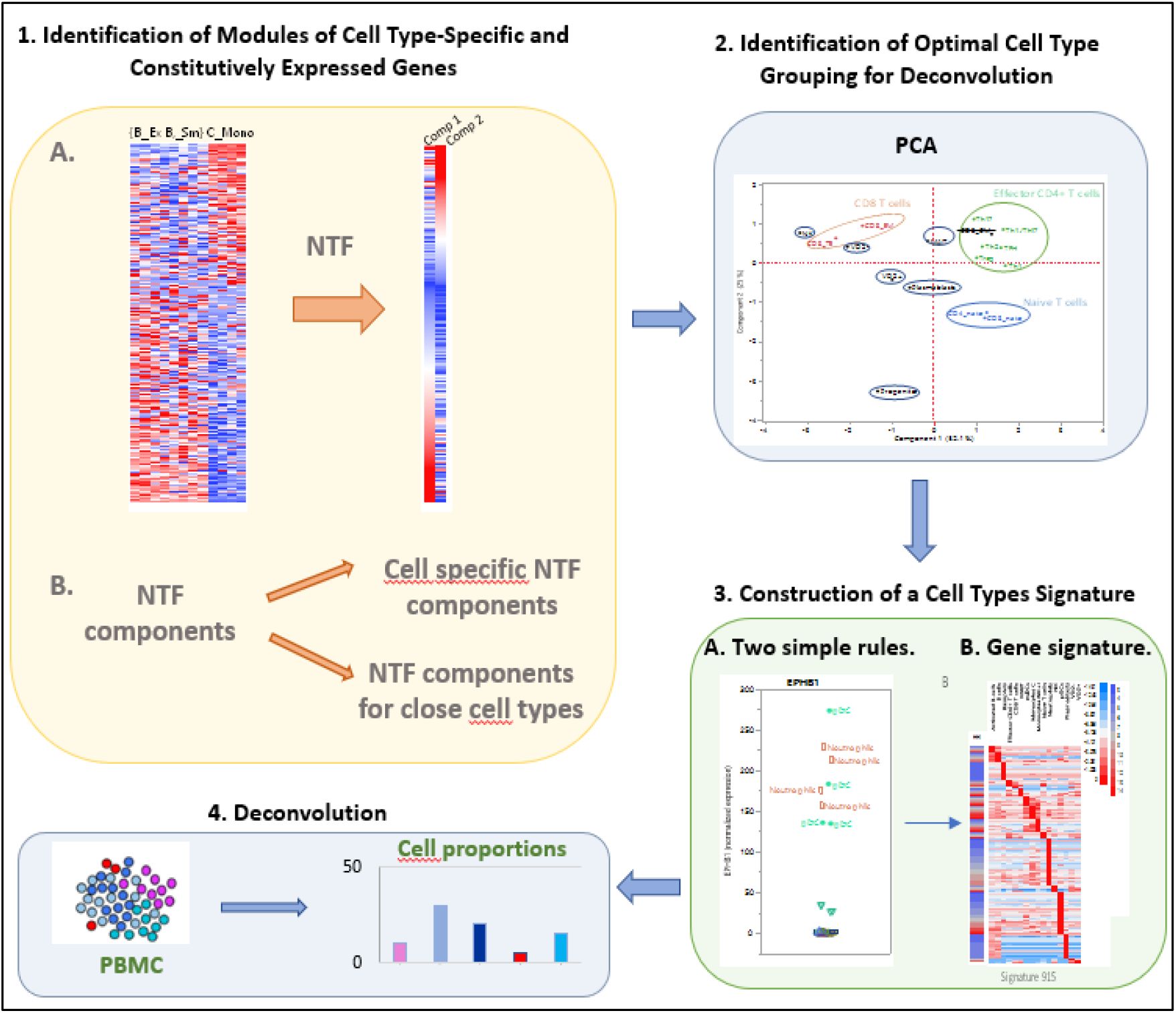

As can be seen from box 1, ASigNTF involves the following steps:

1. Identification of modules of cell type-specific and constitutively expressed genes using NTF
2. Identification of optimal cell type grouping for deconvolution
3. Construction of a cell types signature
4. Deconvolution

Detailed steps are provided in Box 2 and will be discussed in the rest of the article.

### Box 2. Detailed steps to construct highly specific cell type signatures and to perform deconvolution.

**Identification of Modules of Cell Type-Specific and Constitutively Expressed Genes**

- Non-negative Tensor Factorization, NTF, is used for the simultaneous decomposition of expression matrices of subjects from which blood samples were taken and further purified by flow cytometry.
- We use the Herfindahl and Hirschman Index, HHI, to exclude NTF components that are subject-specific.
- Of the other NTF components, the same index allows the identification of two groups. Components from the first group can be specifically associated with unique cell types or pairs of cell types. Components from the second group are used to identify correlated cell types. **Identification of optimal cell type grouping for deconvolution**

- Principal Component Analysis, PCA, is applied to the second group of NTF components.
- The PCA plot allows for identifying closely related cell types.
- The available knowledge can be used to decide whether the grouping of cell types as suggested by the PCA plot is biologically relevant.
- To obtain the gene counts in each identified category, weighted mean counts are calculated, using the flow cytometric proportions as weights for the merged cell types. **Construction of a cell types signature**

- Two rules are applied to identify marker genes:

Rule 1. To represent a specific cell type, a marker gene must have the highest expressions in all samples of that cell type.
Rule 2. To be highly specific for the represented cell type, the HHI of a marker gene should ideally be equal to the number of samples of that cell type.
- In the resulting signature, the median counts observed in the represented cell type of each marker gene are used to rescale counts in the signature matrices.
- The condition number of the scaled signature matrix is calculated. **Deconvolution**

- To estimate the proportions of the cell types in a sample:

1. The median counts observed in the represented cell type of each marker gene are used to rescale the sample counts for the signature marker genes.
2. The vector of scaled counts is projected onto the scaled signature matrix, using non-negative least square regression.
- The PBMC samples from the S13 cohort of the ABIS dataset are used to provide estimates of the RNA abundance factors required to correct regressions estimates for differences in RNA abundance.

### Identification of modules of cell type-specific and constitutively expressed genes using NTF

To illustrate the analytical process, we use as an example a small experiment involving 3 cell types (the data are from the ABIS dataset). It comprises two subtypes of B cells, including Exhausted (B_Ex) and Switched memory B cells (B_SM), and one subtype of monocytes represented by classical monocytes (C_Mono). For each cell type, 4 samples coming from 4 different individuals were profiled by RNA-seq. The counts were normalized for sequencing depth and gene length using TPM method (transcripts per million). The resulting dataset is a concatenation of 4 count matrices – one count matrix per subject. We propose to consider all 4 matrices simultaneously through the use of Non-Negative Tensor Factorization, NTF [13].

The heatmap of the gene expression of the 4 subjects is presented in Figure 1A. The NTF can approximate the gene expression data with only two transcriptional fingerprints, or *gene components*. The 1^st^ gene component (Figure 1B, first column) approximates the gene expression data of the two subtypes of B cells. The 2^nd^ gene component (Figure 1B, second column) approximates the gene expression data of C_Mono. The genes with the highest loadings in the gene components form modules of highly expressed genes.

**Figure 1.**
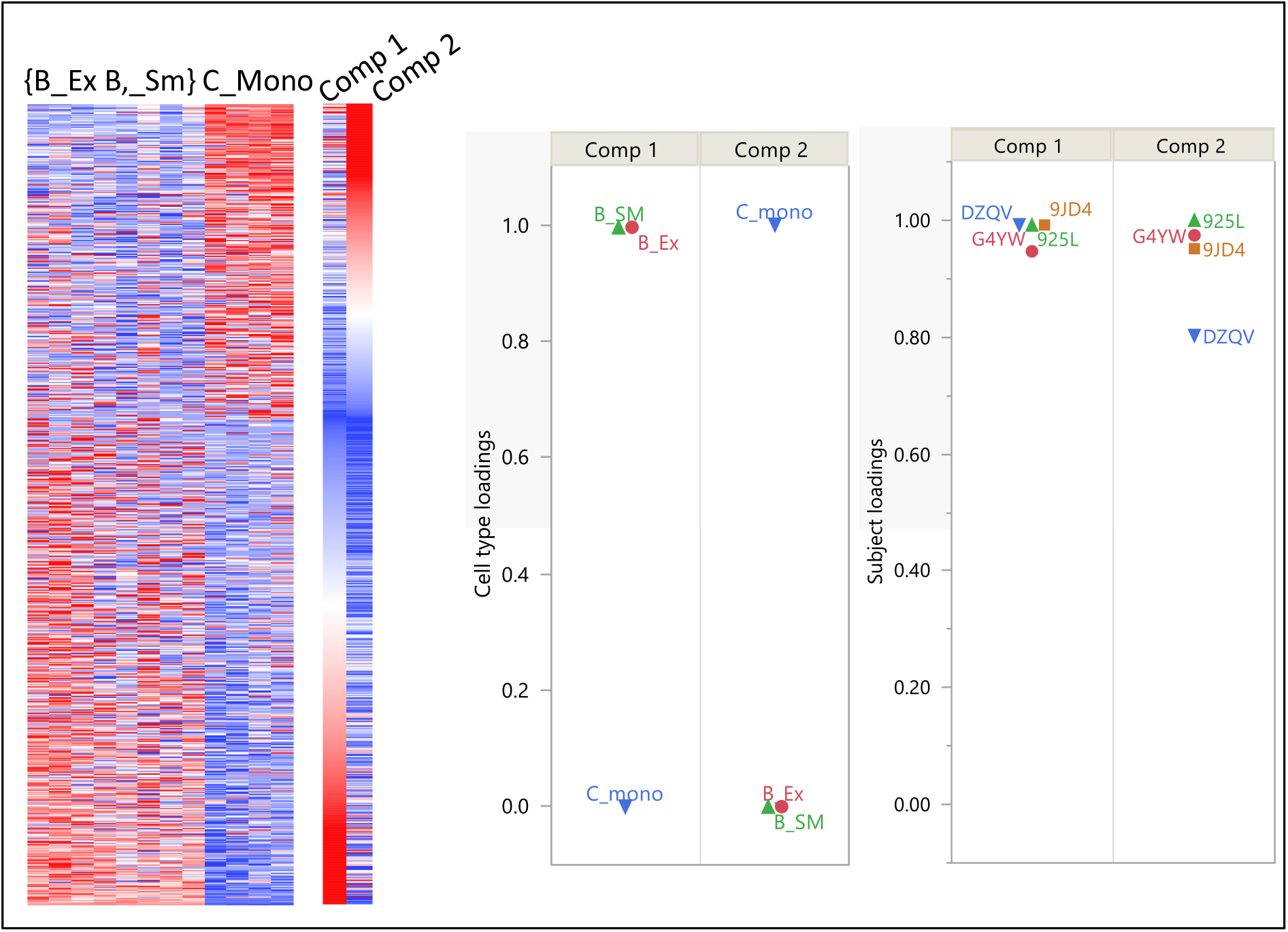
A simple example of the NMF decomposition of three cell types **A**: Expression matrix (red = highly expressed gene, blue = low-expressed gene) of B_Ex, B_SM and C_Mono samples. The first 8 columns in the heatmap correspond to B cells samples and the last 4 columns correspond to classical monocytes samples. **B**: *gene components:* The first component can be associated with B_Ex and B_SM, while the second component can be associated with C_Mono. **C**: cell type and subject loadings: The loadings of B_Ex and B_SM are equal to 1 on the 1st component, while the loading of C_Mono is equal to 0. By contrast, the loading of C_Mono on the 2nd component is equal to 1, while the loadings of B_Ex and B_SM are equal to 0. The 4 subject loadings appear close to 1 on both components, due to the similarities between the 4 subjects.

The NTF also provides the loadings of each cell type on both gene components (Figure 1C left): The loadings of B_Ex and B_SM are equal to 1 on the 1st component, while the loading of C_Mono is equal to 0. By contrast, the loading of C_Mono on the 2^nd^ component is equal to 1, while the loadings of B_Ex and B_SM are equal to 0. These loadings form the corresponding *cell type components*. Although heatmap visualizations of NTF gene components and original gene expression data are intuitively useful to relate gene components to their respective cell types, the same conclusion could be drawn from a post-hoc analysis of the cell type components – and, for this reason, NTF is considered an unsupervised approach.

Furthermore, the specificity of each subject is also considered by estimating the subject loadings on each gene component, forming the correspondent *subject component*. This allows a better approximation to the patterns of gene expression of the individual subjects. The 4 subject loadings appear close to 1 on both components, due to the similarities between the 4 subjects (Figure 1C right).

In summary, the triptych <u>gene</u>, <u>cell type</u>, <u>subject component</u> is called a *component* of the NTF decomposition. The output of the NTF decomposition consists of 3 tables:

1. A table of *cell type loadings* containing as many columns as there are NTF components and as many rows as there are cell types.
2. A table of *gene loadings* containing as many columns as there are NTF components and as many rows as there are genes.
3. A table of *subject loadings* containing as many columns as there are NTF components and as many rows as there are subjects.

Statistical details of the NTF are presented in Box 3.

Like all factorization approaches, NTF is a data reduction technique. The original expression of thousands of genes in a cell type is summarized by the loadings of the cell type on a limited number of gene components, i.e. at the level where a few transcriptional fingerprints, as identified by NTF, are expressed. In our example, this summary could be extended by requiring two additional NTF gene components that would capture the specifics of B_Ex versus B_SM. If more cell types are analyzed by NTF, more gene and corresponding cell type components must be estimated, as in the original ABIS experiment, in which there are 29 cell types (Supp. Table S1).

We now present the NTF analysis of the entire ABIS dataset. One limitation of the NTF is the requirement for a perfectly balanced design, i.e. the same number of genes must be measured for each cell type in each of the 4 subjects. T CD4 Terminal Effector cell type was excluded from the NTF analysis, since data were only available for 2 subjects out of 4. Therefore, 28 cell types were considered for further analysis. Based on the preliminary assumption that the specificity of each cell type can be restored by an appropriate component, 28 components were estimated by NTF. This assumption was further assessed as follows: a component is associated with a specific cell-type if the corresponding cell type loadings are negligible for all cell types except for the one of interest. By contrast, a component is associated with closely related cell types if there are as many elevated loadings in the corresponding cell type component as there are cell types that are close to each other. To characterize the sparsity of a cell type component, we calculated an index derived from the Herfindahl and Hirschman Index [16], which indicates the number of cell types that have non negligible loadings. In our simple example, this index is equal to 2 for the 1^st^ component, since only B_Ex and B_SM have non-zero loading. For the 2^nd^ component, only the classical monocyte cell type has a non-zero loading, so the index is equal to 1. We will further refer to this index as the HHI, and the detailed formula is presented in Box 3.

Similarly, the HHI can be calculated for the corresponding subject loadings. If the NTF component is shared uniformly by all subjects, then the HHI is equal to the number of subjects; conversely, if only one subject is associated with that component, then the HHI is equal to 1.

For each NTF component, the sparsity of the corresponding cell type component was analyzed to assess their cell type specificity. The sparsity of the corresponding subject component was also evaluated to determine if the NTF component is shared uniformly by most subjects. Only those components that are common to most subjects are of interest since we are not interested in the specificities of each subject. Other components are not considered further.

The results are shown in the scatterplot depicting the HHI of the 28 NTF components. For each component, the HHI is calculated for both: the subject component (Y-axis) and cell component (X-axis) loadings (Figure 2). Each point corresponds to a component: 15 of the 28 components in the lower part of the figure have a subject component HHI that is lower or close to 2 (grey zone), indicating that these components are shared by at most 2 out of 4 subjects and can be ignored due to their subject specificity. The 13 left components in the upper part of the figure have a subject component HHI higher or close to 3, indicating that these components are shared by at least 3 out of 4 subjects. Two groups can be identified from these 13 NTF components: The first group of 8 NTF components on the left side (red, solid colored circles) has an HHI that is lower or close to 3, indicating that these components are shared by at most 3 cell types. These components are considered cell type-specific.

**Figure 2.**
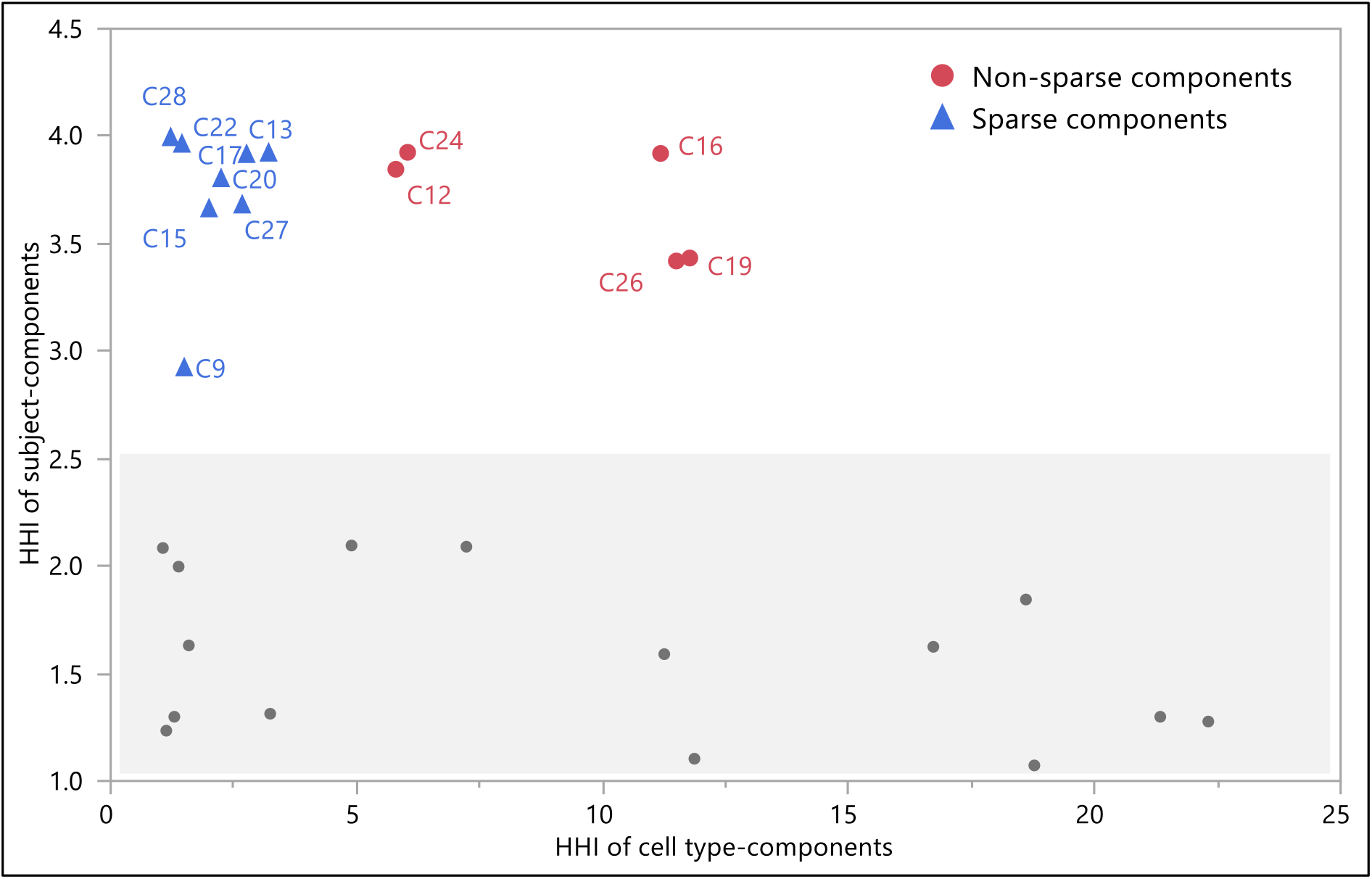
Scatterplot of the HHI of the 28 subject components (Y-axis) compared to the HHI of the corresponding cell type components (X-axis). Each point corresponds to a component: 15 of the 28 components in the lower part of the figure have a subject component HHI that is lower or close to 2 (grey zone), indicating that these components are shared by at most 2 out of 4 subjects. The 13 left components in the upper part of the figure have a subject component HHI higher or close to 3, indicating that these components are shared by at least 3 out of 4 subjects. Two groups can be identified from these 13 NTF components: The first group of 8 NTF components on the left side (blue triangles) has an HHI that is lower or close to 3, indicating that these components are shared by at most 3 cell types. These components are considered cell type-specific. The other 5 NTF components on the right side of the figure (red, solid colored circles) have an HHI greater than 5, indicating that these components are shared by at least 5 cell types. These components form a second group and are nonspecific, since they are common to a larger number of cell types.

The other 5 NTF components on the right side of the figure (blue crosses) have an HHI greater than 5, indicating that these components are shared by at least 5 cell types. These components form a second group and are considered to be nonspecific. All in all, we find that by estimating a maximum number of components and later ignoring irrelevant components based on clearly defined criteria, we overcome the classic factorization problem of pre-selecting a number of components whose optimality depends on the mathematical criterion used.

### Identification of optimal cell type grouping for deconvolution

For 8 specific NTF components coming from the previous step of analysis the distributions of the cell type loadings are presented in Figure 3, using abbreviated names for cell types (Supp. Table S1). From these distributions, we see that components 9, 13, 20, 22 and 28 are specific to Neutrophils, Myeloid Dendritic cells (mDC), Classical monocytes (C_Mono), Plasmacytoid Dendritic cells (pDC) and Basophils, respectively. From the 3 other components, pairs of close cell-types can be identified: B_Ex and B_SM on component 15, Non-classical monocytes (NC_mono) and Intermediate monocytes (I_mono) on component 17, and Non-switched B memory cells (B_NSM) and Naïve B cells (B_Naive) on component 27.

**Figure 3.**
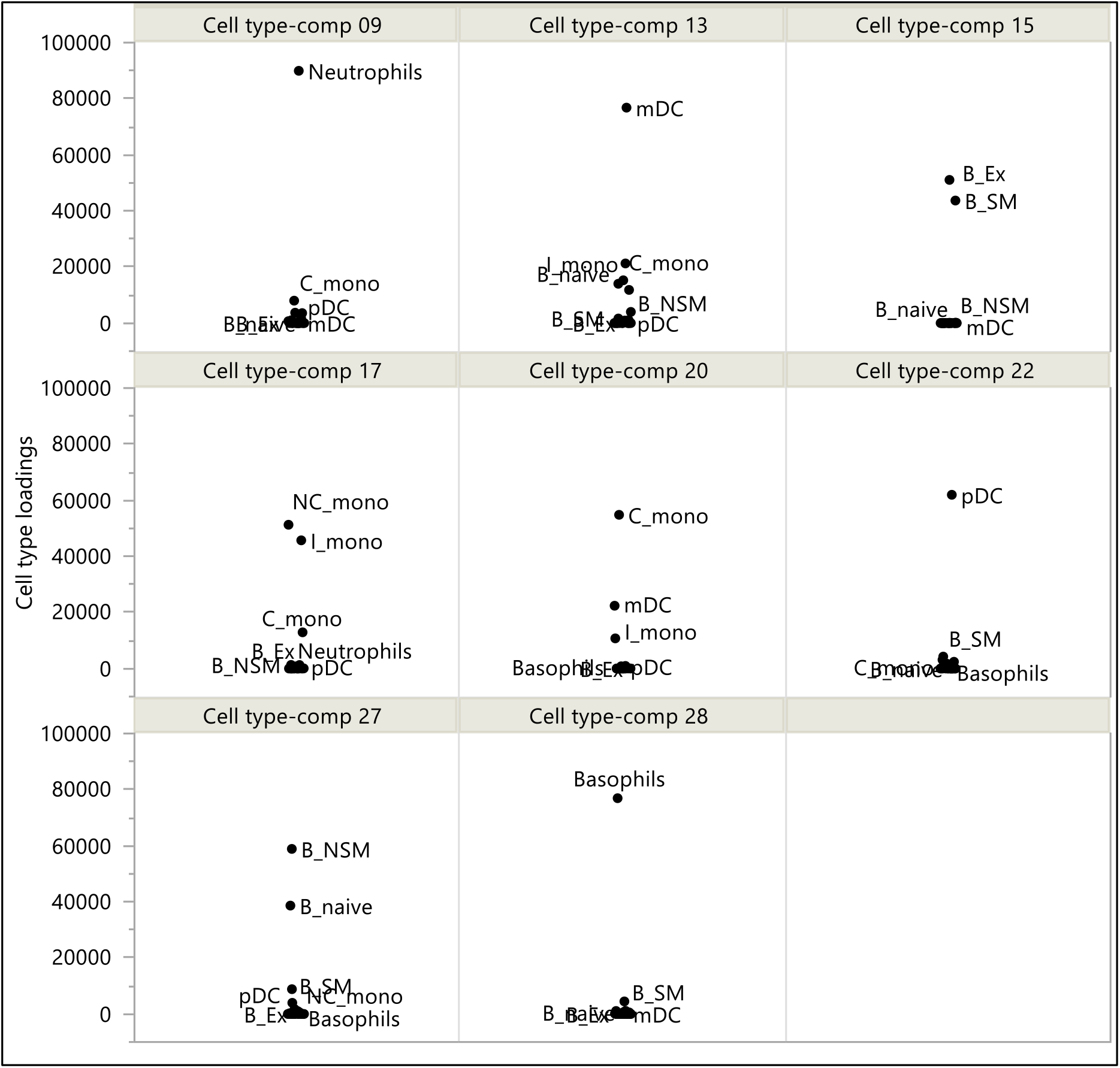
Cell types coming out with the highest loadings for 8 NTF sparse components. From these distributions, we see that Neutrophils, mDC, C_Mono, pDC and Basophils load exclusively on their respective gene components and are therefore specific. Two pairs of close cell-types can be identified: B_Ex and B_SM from the gene component 15, and B_NSM and B_Naive from gene component 27.

Deconvolution only works for cell types that exhibit a detectable and specific signal pattern. Cell types with remarkably similar transcriptomic profiles need to be grouped. To define the grouping of cell types, we have used an agnostic, unsupervised method, which consists in performing a Principal Component Analysis, PCA, on the 5 nonspecific cell type components (Figure 4). The 11 cell types that were associated with specific NTF components, form a separate cluster (Supp. Figure S1). They were excluded from the PCA to focus on the similarities between the remaining cell types.

**Figure 4.**
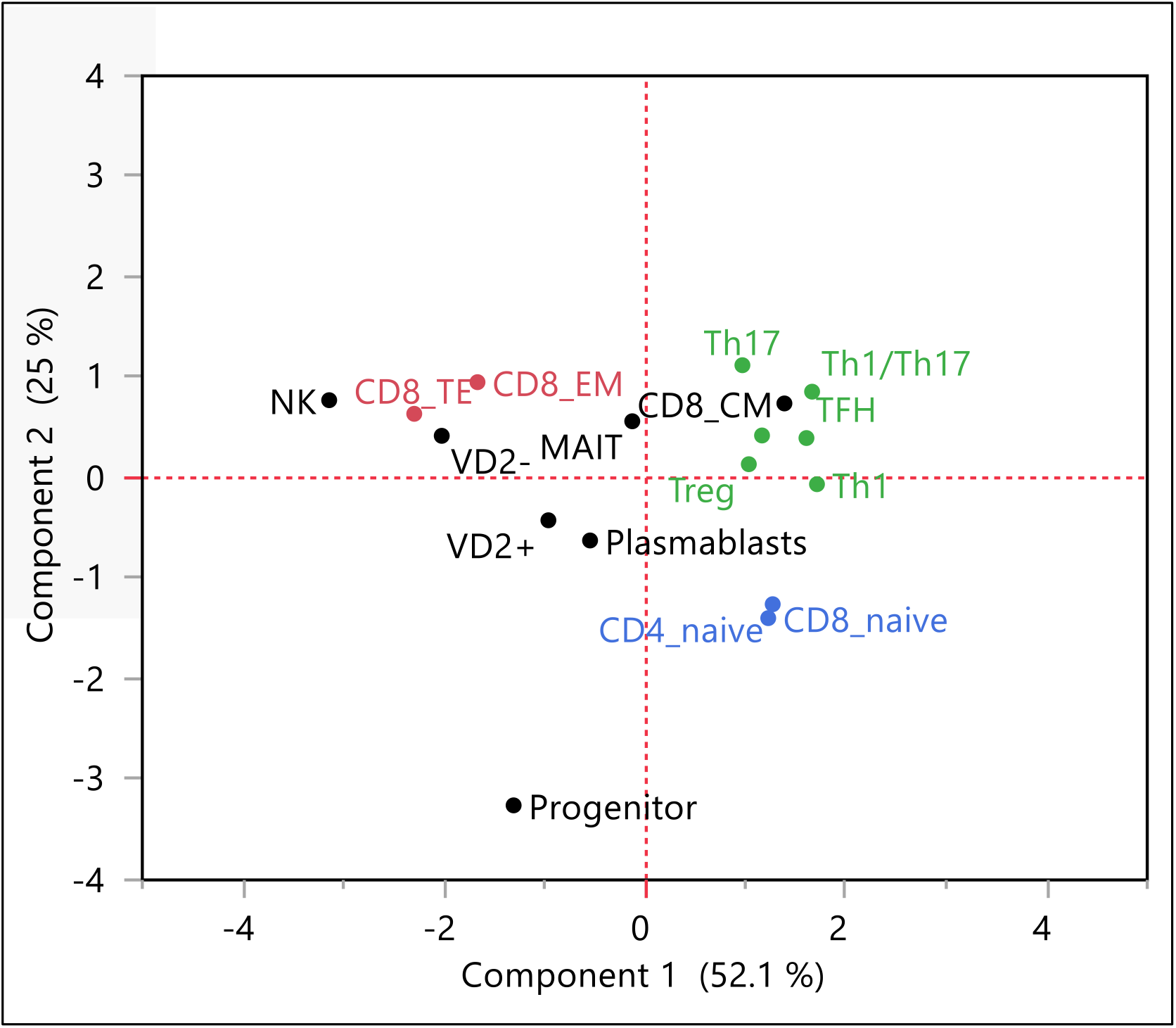
A Principal Component Analysis was applied to the 5 nonspecific cell type components to define the grouping of cell types with close transcriptomic profiles. Three subgroups for **CD8 T cells** (CD8_TE and CD8), **Effector CD4+ T cells** (Th17, Th1/Th17, Th2, TFH, Treg, Th1) and **Naive T cells** (CD4_naive and CD8_naive) were defined for closely related cell types: NK and Gamma-delta VD2-T cells were considered separately from CD8 T cells. MAIT, Plasmablasts and Progenitor are considered as distinct cell types. Gamma-delta VD2+ and VD2-T cells are considered as separate classes, being too far apart to be regrouped. CD8_CM is considered a biological outlier and not grouped with effector CD4+ T cells.

For cell types from the two main classes CD4+ and CD8+ T cells, the PCA (Figure 4) allowed the definition of 3 subgroups of closely related cell types: CD8+ T cells (CD8_TE and CD8_EM), effector CD4+ T cells (Th17, 1h1/Th17, Th2, TFH, Treg and Th1) and naive T cells (CD4_naive and CD8_naive). Both CD4+ and CD8+ cells can have different states, including naive, effector or memory phenotypes. Naïve T cells of both classes were found to be transcriptionally close. This difference was more pronounced after cells priming, resulting in a clear separation of cell types belonging to the effector CD4+ and CD8+ T subgroups. Thus, our agnostic, non-supervised method shows that CD4_naive and CD8_naive have tightly similar transcriptomic fingerprints and can be grouped together, despite the widely accepted classification in two different cell types.

NK and Gamma-delta VD2-T cells must be considered separately as they are biologically different from CD8 T cells. NK cells, as well as CD8+ T cells, have cytotoxic functions, which may explain their proximity in transcriptomic profiles, as PCA has shown. However, NK cells are considered separately as they represent a separate class of lymphocytes. MAIT, Plasmablasts and Progenitor cell types appear as distinct cell types. Gamma-delta VD2+ and VD2-T cells are too far apart to be regrouped. CD8_CM is considered a biological outlier since it is grouped with effector CD4+ T cells instead of being close to CD8 T cells. Therefore, this cell type was not used for deconvolution. The progenitor cell type was further ignored given its very low concentration in known PBMC samples, making it unsuitable for deconvolution.

To emphasize the crucial role of NTF cell type components, we also performed the PCA on the matrix of normalized counts with 28 cell types as rows and 4 x p genes as columns, where p is the number of genes. The analysis process comparing both PCA analyses is summarized in Figure 5.

**Figure 5.**
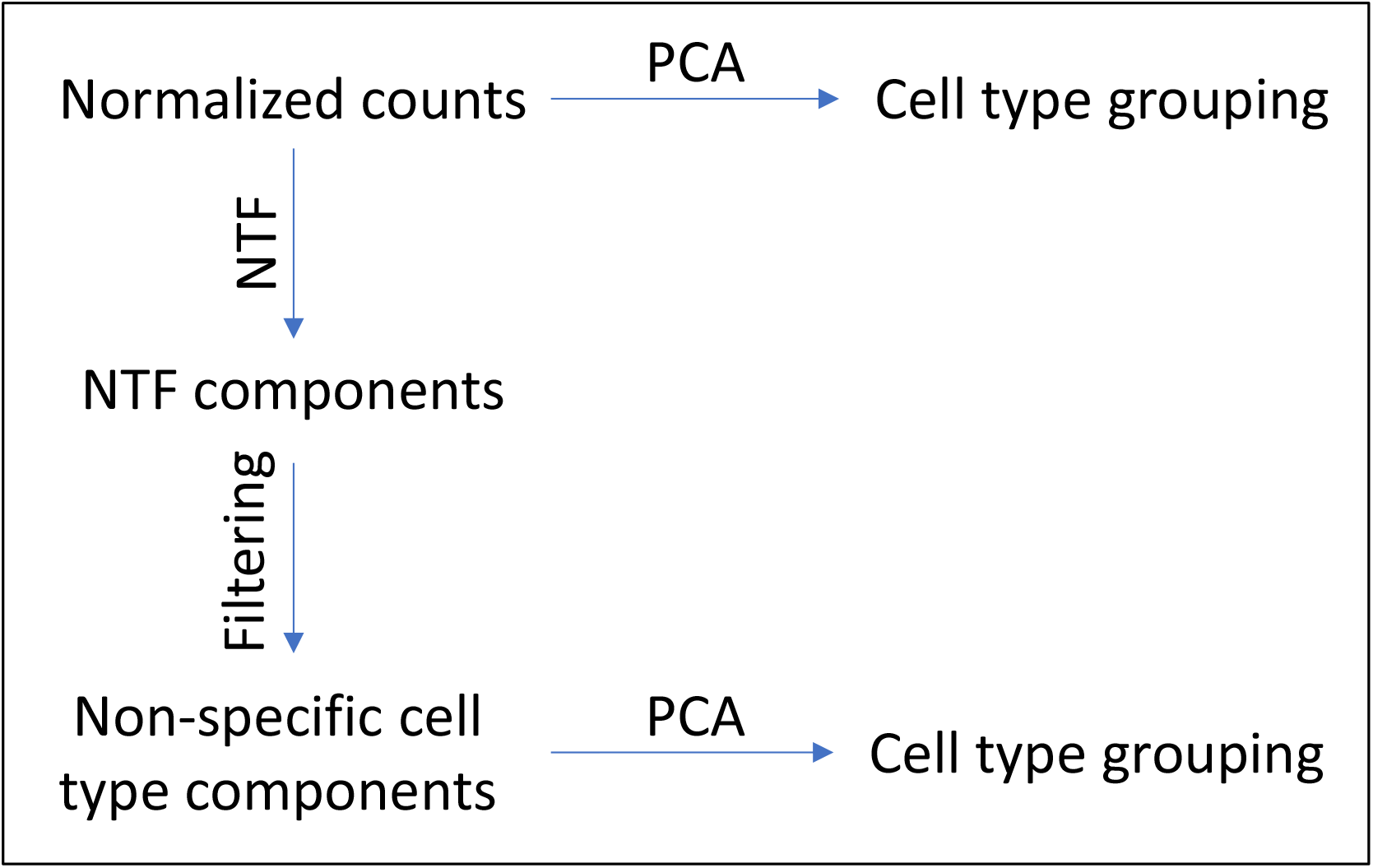
Scheme of the PCA analysis process on normalized counts versus NTF nonspecific components.

In contrast to the PCA analysis based on NTF cell type loadings, only large groups could be identified (Figure 6). In fact, 6 possible cell groups, comprising Neutrophils, Plasmablasts, Progenitors, Myeloid lineage cell group, group of B and T cells, can be identified on this PCA (Figure 6). Unlike the PCA performed on NTF components, there is no separation between two major classes of T cells, including CD4+ and CD8+ T cells. Our NTF-based processing of the data enables a more efficient extraction of the information needed to define closely related cell types. First, components corresponding to transcriptomic variations that are subject-specific are ignored. Secondly, cell types that can be uniquely defined by specific components are excluded.

**Figure 6.**
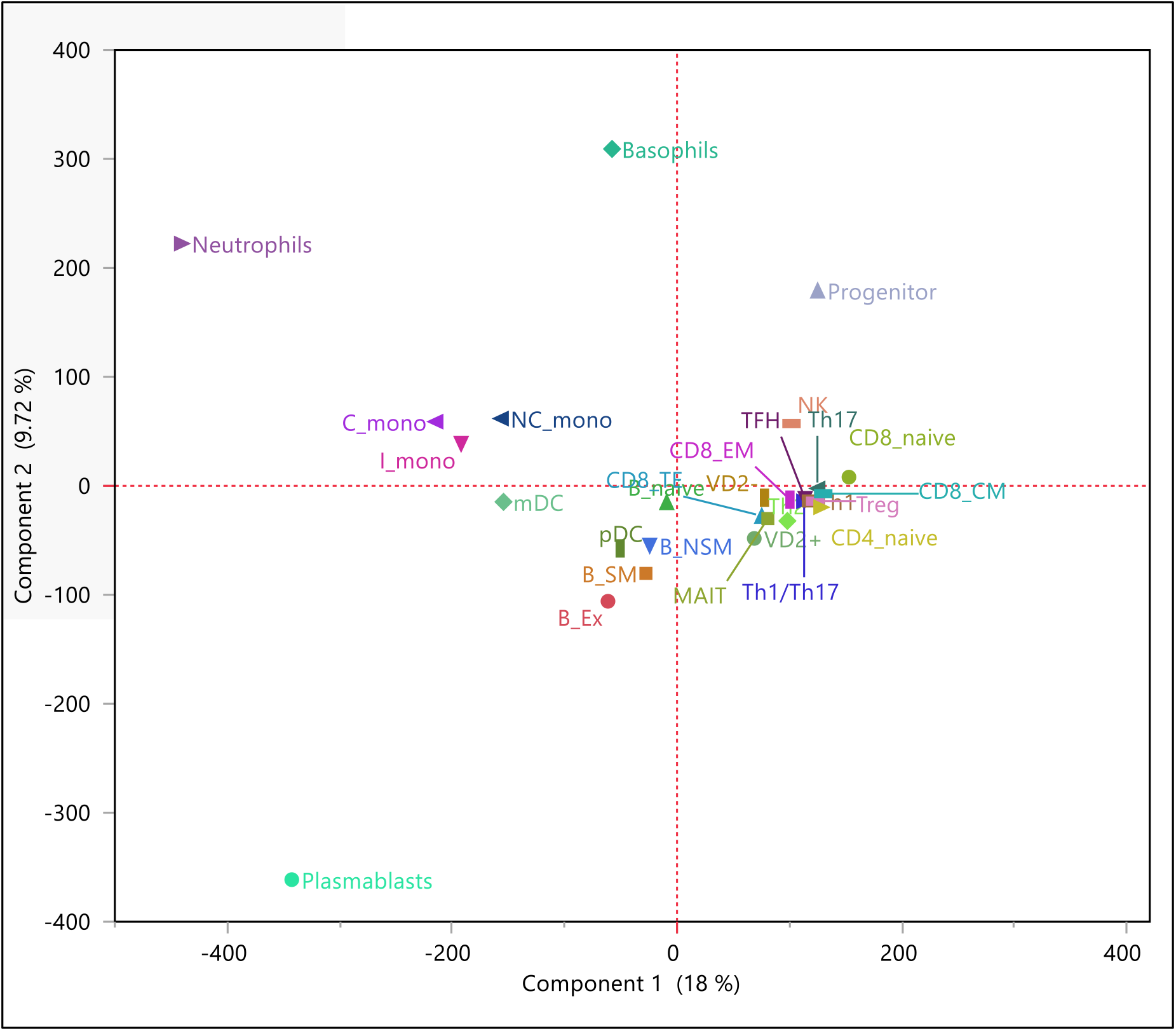
PCA on the matrix of normalized counts with 28 cell types as rows and 4 x p genes as columns, where p is the number of genes. Unlike the PCA performed on NTF components, there is no separation between two major classes of T cells, including CD4+ and CD8+ T cells.

In total, 16 categories of cell types, including either specific or regrouped cell types, were identified (Table 1A). Of note, these 16 categories are very close to the 17 categories that were proposed by Monaco et al [1] (Table 1B). However, our approach is totally unsupervised, while the original approach attempted numerous cell types combinations to maximize the correlation results between true and estimated proportions in left-out PBMC samples that were submitted to deconvolution. Of course, we still used our biological knowledge to decide whether the grouping of cell types that are close to each other in our data-driven analysis is relevant. To obtain the gene counts in each identified category, we calculated weighted mean counts, using the flow cytometric proportions as weights for the merged cell types.

**Table 1.** Categories of cell types, NTF and ABIS approaches

| Category NTF | Cell type components | Category ABIS | Cell type components |
| --- | --- | --- | --- |
| Activated B cells | B_Ex + B_SM | B Memory | B_NSM, B_SM, B_Ex |
| B cells | B_NSM + B_Naive | B Naïve | B_Naive |
| Basophils | Basophils | Basophils | Basophils |
| Effector CD4+ T cells | Th1/Th17, Th17, Th2, Th1, TFH, Treg | T CD4 Memory | Th1/Th17, Th17, Th2, Th1, TFH, Treg, T CD4-TE |
| CD8 T cells | T CD8_TE, T CD8_EM | T CD8 Memory | T CD8_TE, T CD8_EM, T CD8_CM |
| MAIT | MAIT | MAIT | MAIT |
| mDCs | mDCs | mDCs | mDCs |
| Monocytes C | Monocytes C | Monocytes C | Monocytes C |
| Monocytes NC+I | NC_Mono + I_Mono | Monocytes NC+I | NC_Mono + I_Mono |
| Naïve T cells | T CD4_naive, T CD8_naive | T CD4 Naïve | T CD4_naive |
| Neutrophils | Neutrophils | T CD8 Naïve | T CD8_naive |
| NK | NK | Neutrophils | Neutrophils |
| pDCs | pDCs | NK | NK |
| Plasmablasts | Plasmablasts | pDCs | pDCs |
| VD2- | VD2- | Plasmablasts | Plasmablasts |
| VD2+ | VD2+ | VD2- | VD2- |
|  |  | VD2+ | VD2+ |
| **A** |  | **B** |  |

### Construction of a cell types signature

Once the cell types have been regrouped, it is possible to identify marker genes of each individual cell type or group of cell types (for simplicity, in what follows, the 16 identified categories will be referred to as cell types). A traditional approach is to compare each cell type with the pool of all other cell types [1] and to retain the most differentially expressed genes with a fold change exceeding an arbitrary cutoff. As the following example illustrates, this approach does not necessarily identify clear marker genes.

Performing differential analysis, EPHB1 and LIN7A can be identified as marker genes of Neutrophils (Figure 7A), based on extremely significant p-values and high folds (p = 8.e-9; FDR-adjusted [17] p = 8.e-8; fold = 460, and p = 4.e-8; FDR-adjusted p = 4.e-7; fold = 58, respectively). However, EPHB1 is also a marker gene of pDC, while LIN7A is also a marker gene of Basophils. In contrast, KRT23 and KCNJ15 are exclusive marker genes of Neutrophils (Figure 7B).

**Figure 7.**
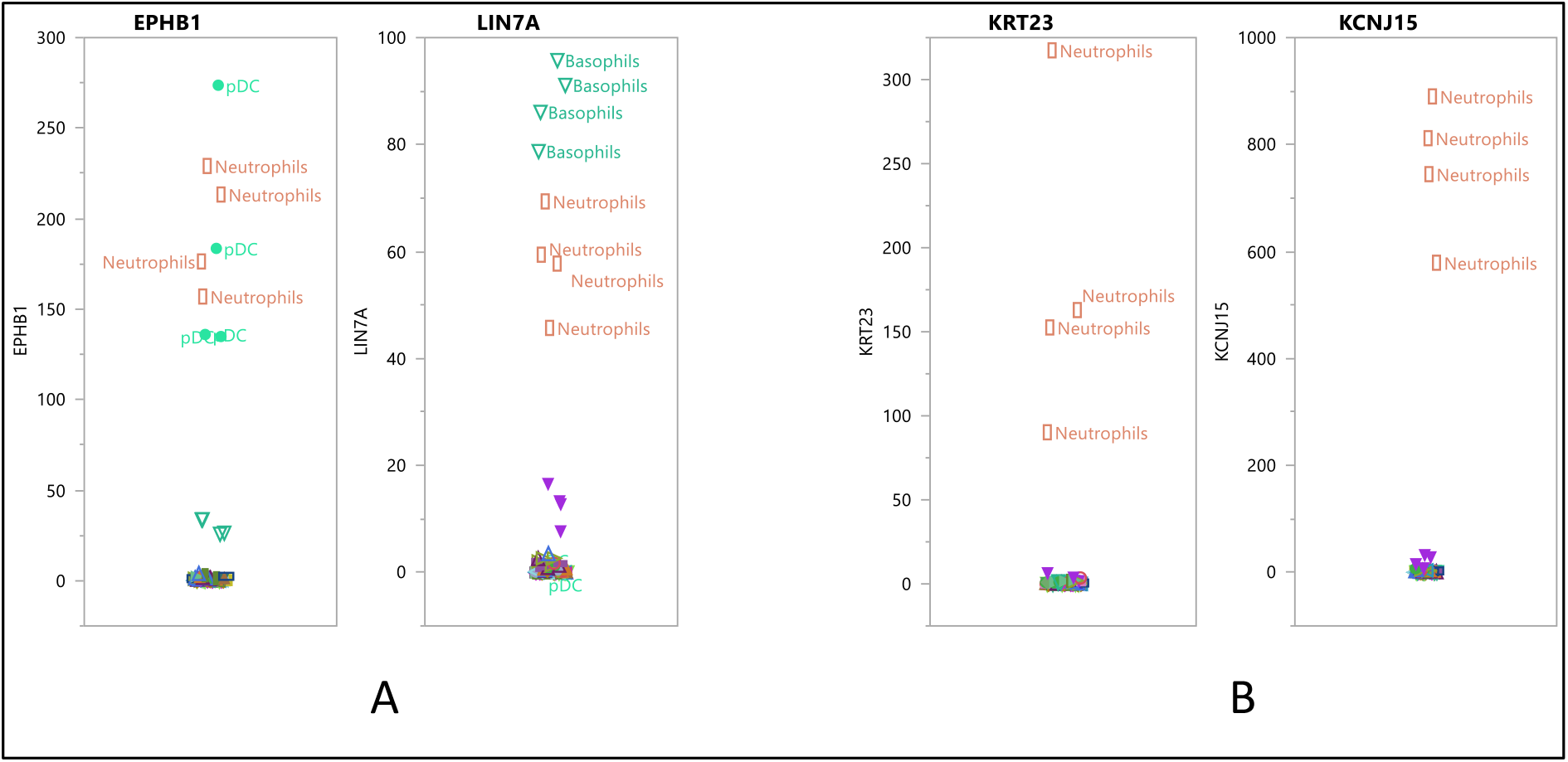
Exclusive and non-exclusive Marker Genes **A:** EPHB1 and LIN7A are marker genes of Neutrophils, in terms of p-values and folds (p = 8.e-9; fold = 460 and p = 4.e-8; fold = 58, respectively). EPHB1 is also a marker gene of pDC while LIN7A is also a marker gene of Basophils. **B**: In contrast, KRT23 and KCNJ15 are exclusive marker genes of Neutrophils.

To prevent these situations, we propose using the following two simple rules:

Rule 1: To represent a specific cell type, a marker gene must have the highest expressions in all samples of that cell type.
Rule 2: In order to be highly specific for the represented cell type, the HHI of a marker gene should ideally be equal to the number of samples of that cell type.

Going back to our examples, Rule 1 would exclude EPHB1 and Rule 2 would exclude LIN7A, because HHI is higher than 11. Both KRT23 and KCNJ15 satisfy the two rules, with an HHI equal to 4. It is remarkable that the first rule is supervised by the cell type, while the second rule is unsupervised.

To validate our signatures on datasets coming from both microarray and RNA-seq experiments, we first filtered out genes detected in the ABIS-seq experiment, which are not covered by probesets in microarrays. For that, we considered the intersection between ABIS-seq transcripts and the microarray dataset GSE65133 [15], resulting in a subset of 14137 genes. For practical consideration, only genes with a median TPM-normalized count higher than 10 in their represented cell type were evaluated to ensure that selected marker genes are sufficiently expressed to be detected in mixtures. The gene number passing the first rule ranges from 7 in plasmablasts to 112 in neutrophil LD, which leads to an over-representation of the latter. The gene number passing the second rule ranges from 1 in NK to 30 genes in basophils. Therefore, the HHI value used in the second rule must be relaxed for cell types with under-represented marker genes numbers. The relaxation of the HHI occurs at the expense of marker gene specificity in under-represented cell types and raises a problem known as multicollinearity. Therefore, the number of marker genes should be reduced for over-represented cell types to avoid masking or suppression of less represented and potentially correlated cell types [18]. In practice, genes that fulfill the first rule are ranked by cell type and increasing HHI value. Further, up to 30 marker genes per cell type are selected. This procedure yields a signature of 415 marker gene of 14 cell types – VD2+ and VD2-were excluded since no marker genes satisfied the first rule. In the signature heatmap (Figure 8A), each row corresponds to a marker gene and each column corresponds to the counts of the marker genes in a given cell type.

**Figure 8.**
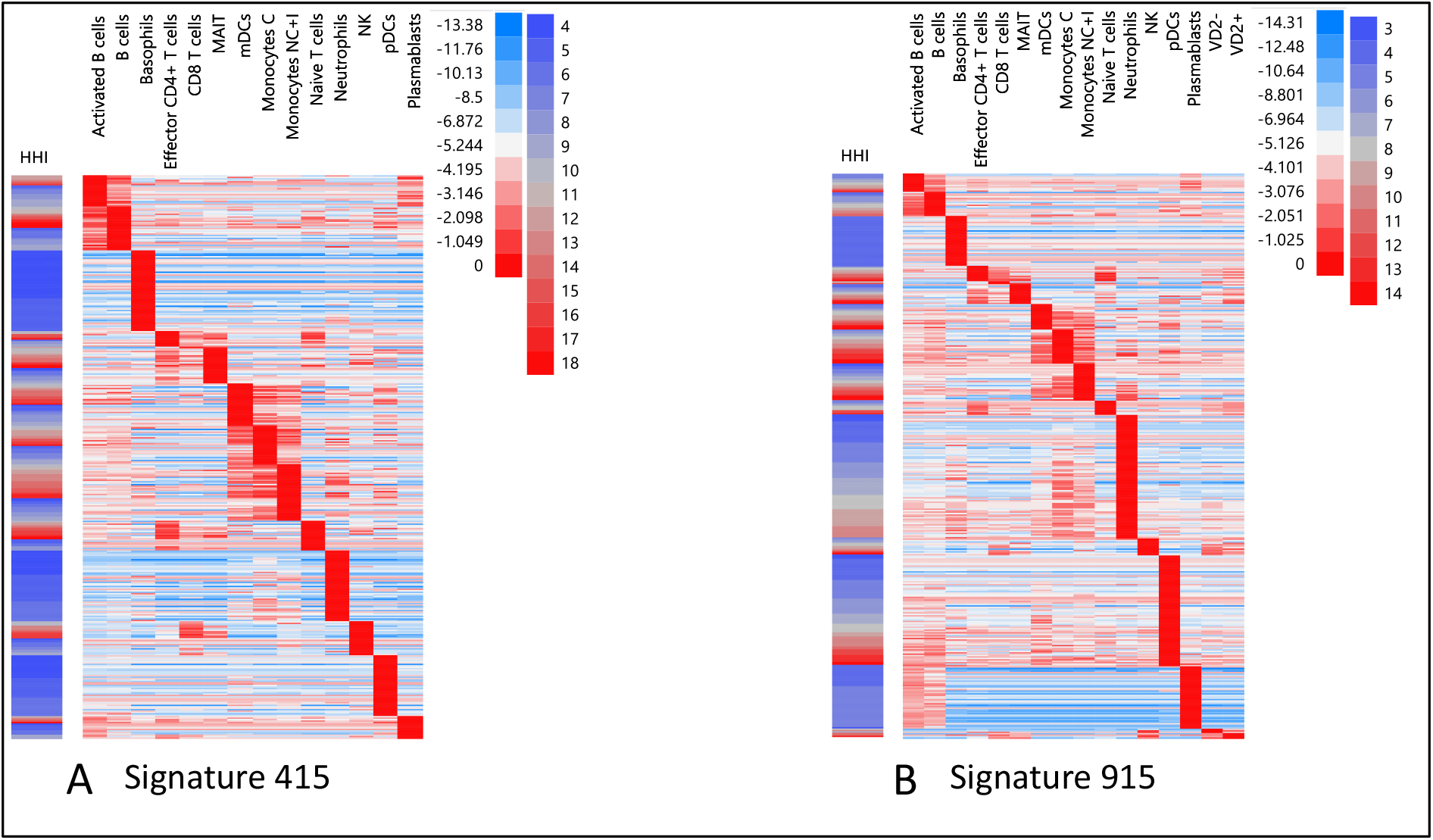
Heatmaps of 415 and 915 genes signatures. In the signature heatmaps, each row corresponds to a marker gene and each column corresponds to the log2-expression of the marker genes rescaled by the median count observed in the represented cell type. **A**: Among a subset of the transcripts identified in the ABIS-seq experiment, which are also commonly found in microarrays, genes with the highest expressions in *all* samples from the same cell type are identified as marker genes. For each cell type, marker genes were sorted by ascending HHI and a cutoff was chosen that *ensures at most 30 marker genes*. **B**: Same procedure as in A using all genes measured in the ABIS-seq experiment, with an objective of *at most 50 marker genes by cell type* and allowing genes with the highest expressions in 3 out of the 4 samples from the same cell type if the objective of *4 out of 4 samples* cannot be met.

We also propose a less restrictive procedure, *allowing at least 3 out of the 4 samples of the same type to be on top of the counts –* if the objective of 4 samples cannot be met – and looking at all genes listed in the ABIS experiment (37786). Given the substantially higher number of genes, more candidate marker genes can be discovered, so the maximum allowable number of genes per cell type can be increased to 50. This procedure yields a signature of 915 marker genes of all 16 cell types, this time including VD2+ and VD2-(Figure 8B).

As will be seen in the next section, our deconvolution algorithm uses a non-negative least square regression [14]. Due to the skewed distribution of counts, the regression discrepancies tend to be smaller at low counts, so that less expressed marker genes tend to have less influence. In order to give equal importance to low and high expressed genes, the median counts observed in the represented cell type of each marker gene are used to rescale counts in the signature matrices – and in the sample to be deconvoluted to ensure consistency. Following this rescaling, the median count of a marker gene in the represented cell type is always 1.

The exclusive character of the discovered marker genes for most of the cell types is visually clear on both heatmaps. The almost exclusive expression in a particular cell type is reflected by the red segments on the diagonal of the heatmap, with the exception of a few cell types where the rules have been relaxed to ensure a sufficient number of marker genes per cell type (e.g. Effector CD4+ T cells and CD8 T cells, or VD2+ and VD2-in the signature of 915 genes). The specificity of marker genes for a particular cell type can be fully confirmed by calculating the condition number of the signature matrices, which should be as small as possible to ensure a low degree of multicollinearity between cell type signatures. Signature 415 has a kappa of 13.1 and signature 915 has a kappa of 7.2, to be compared with CIBERSORT’s LM22 signature [15] (kappa = 11.3) and ABIS-seq [1] (kappa = 12.9). Thus, our signature 915 surpasses CIBERSORT’s LM22 signature with a 36% reduction in the condition number achieved while our signature 415 is comparable with the ABIS-seq signature. Statistical details on the calculation of the condition number are presented in Box 3.

#### Box. 3 Statistical details

**Non-Negative Tensor Factorization**

In the ABIS experiment, four PBMC samples from four different subjects were subjected to flow cytometric sorting. Each filtrate corresponds to a specific cell type and it was further profiled by RNA-seq. The counts were normalized for sequencing depth and gene length using TPM method (transcripts per million).

The resulting dataset is a longitudinal concatenation of four count matrices **X_i_** (n x p) – one count matrix per subject. We propose to consider all four matrices simultaneously through the use of Non-Negative Tensor Factorization, NTF [Cichocki], which extends the NMF to longitudinal data as follows: NTF estimates cell type components and gene components common to all subjects. Nevertheless, each subject may be more or less associated with a particular gene component. The specificity of each subject is taken into account during the estimation process by an additional vector containing the loadings of each subject on the gene component of interest. The result is still an additive mixture, with the following formula:

**X = W** ⊗ **H** ⊗ **Q**, where ⊗ denotes the tensor product.

- **X**(*n* x *p* x *q*) is a matrix of counts, where *n* is the number of cell types, *p* is the number of genes and q is the number of subjects.
- The columns of **W** (*n* x *k*), **H** (*p* x *k*) and **Q** (*q* x *k*) are the cell type components, gene components and subject components, respectively.
- *k* is the number of gene components and corresponding cell type components.

**W**, **H** and **Q** are constrained to contain only non-negative entries, ensuring the interpretability of the columns of **H** as gene components and their property of additivity – through the non-negative columns of **W** and **Q** – to approximate real samples.

**W**, **H** and **Q** can be estimated by the HALS algorithm [Cichocki].

**Herfindhal-Hirschman Index, HHI**

The index **HHI** is mathematically defined by the following formula:

HHI = Round (Sum(*x*_i_)^2^ / Sum(*x*_i_^2^))

where the *X*_i_’s are the elements of the vector for which we want to evaluate the sparsity.

The HHI varies between 1 and *n*, where *n* is the size of the vector. It indicates the number of entries in the vector that are not negligible.

**Condition Number**

To calculate the condition number, Singular Value Decomposition, SVD, is performed on the signature matrix. The condition number, or *kappa*, can be defined as the ratio of the first singular value to the last singular value. The closer the Kappa is to 1, the more uncorrelated are the cell types of the signature matrix.

### Deconvolution

To estimate the proportions of the cell types in a sample, the vector of gene counts is projected onto the signature matrix. Several methods have been proposed, e.g. robust linear regression [1], support vector regression with linear kernel, ν-SVR [15], non-negative least square regression, nnls [2] [19]. The robust linear regression implements an unconstrained optimization problem so that negative proportions may arise [1]. ν-SVR is known to address the problem of multicollinearity [20] [21]. Our approach provides highly specific signatures as shown by their condition number, reducing the effects of multicollinearity. This might explain why the application of nnls to the PBMC samples from the S13 cohort of the ABIS dataset, as described in the next section, worked as well as other methods, which motivated us to continue using nnls because of its ease of use. It is noteworthy that we have not evaluated other methods against other benchmark data to ensure a comprehensive comparison. Thanks to the non-negativity granted to them, regression estimates can be further normalized by their sum to meet the ‘sum-to-1’ condition – as expected when it comes to proportions. This additional normalization may not be applied because contents unknown to the applied signature can lead to absolute estimates for immune cell types that do not add up to 1.

The signature is based on normalized RNA-seq counts from purified filtrates, where different gene counts reflect the differences in expression *within* a given cell type. However, in a bulk sample, e.g. whole blood or PBMC, a cell type that contains more RNA will also contribute more to the total gene count than a cell type that contains less RNA, even if they are in equal proportions – a fact that is masked when purified filtrates are considered. Therefore, correction factors must be applied to the regression estimates to correct for differences in RNA abundance [1]. The PBMC samples from the S13 cohort of the ABIS dataset were used to provide estimates of the RNA abundance factors. Specifically, for each cell type the estimates found by nnls were regressed on the true proportions as found by flow cytometry.

Since RNA-seq counts were used to define our signatures, target quantiles are needed to normalize microarray gene expressions. These target quantiles are based on the flow cytometric samples from the ABIS experiment.

### Estimating mixture proportions in benchmark expression data

We evaluate our results using several public datasets focused on PBMC and whole blood tissues and compare them with those obtained by available signatures and software, namely ABIS-seq, ABIS-microarray and CIBERSORT [1] [15].

The Venn diagram shows a small degree of overlap between our signatures and the two external ABIS-seq and CIBERSORT signatures (Figure 9), which illustrates the specific features of our signatures. The CIBERSORT signature shares the smallest number of genes with the other 3 signatures.

**Figure 9.**
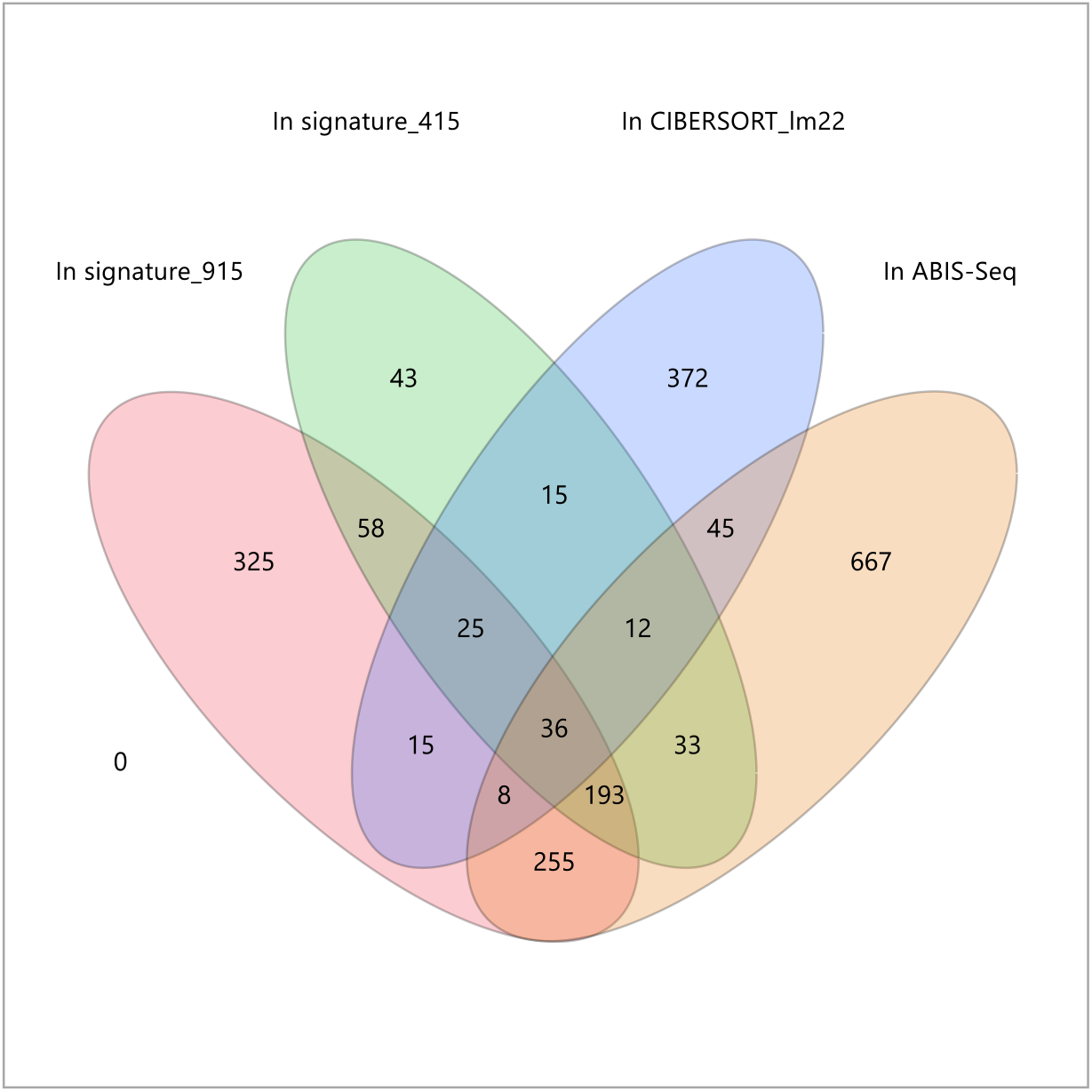
Venn diagram between signatures 415 and 915, ABIS-seq and CIBERSORT. The CIBERSORT signature shares the least number of genes with the other 3 signatures.

Four datasets with diverse contents were used to benchmark the signatures:

- Deconvolution of purified immune cell types
- Immune cell types proportions from PBMC samples profiled by RNA-seq
- Mixtures with unknown content
- Immune cell types proportions from PBMC samples profiled by microarray

### Deconvolution of purified immune cell types

Before dealing with cell type mixtures, we first evaluated the performance of each deconvolution method in identifying isolated cell types. We used the GSE118165 study [22] in which 25 immune cell types, including various subgroups of B cells, CD4+ T cells, CD8+ T cells, Gamma-delta T cells, monocytes, dendritic cells (DCs) and natural killer cells (NK) were isolated by flow cytometry from peripheral blood of up to four healthy donors. Sorted immune cells were activated and subjected to RNA-seq. Following pseudoalignment, gene abundances were estimated using tximport package. For each signature, as shown in Figure 10 using NK cells as an example, the 3 cell types with the highest estimated proportions were considered. In the particular case of the ABIS-seq package, we found that the proportion for NK cells was higher than 100% and that some proportion estimates were negative due to the absence of a non-negative constraint for the robust regression used. Therefore, it was not possible to perform the sum-to-1 normalization required to calculate the proportion estimates as was the case with our application and CIBERSORT.

**Figure 10.**
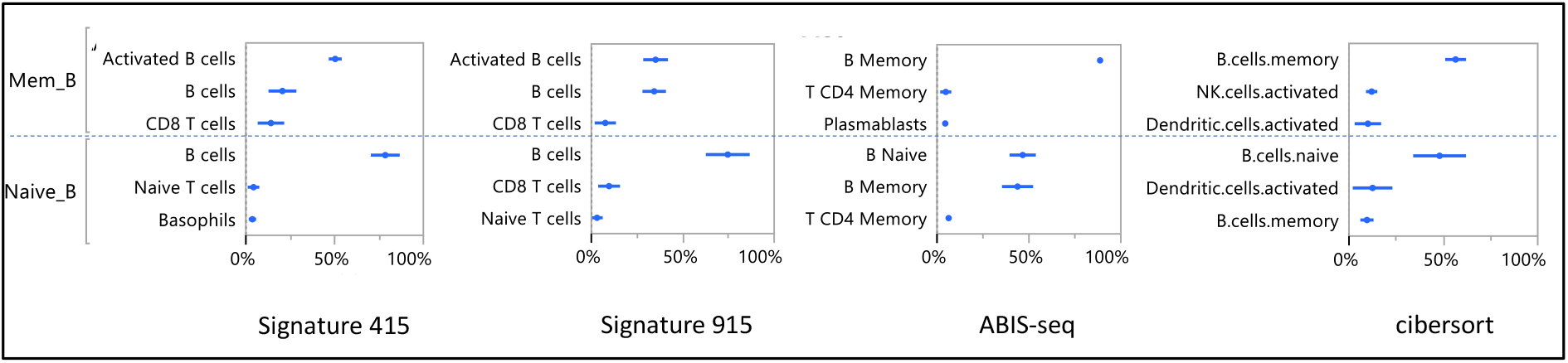
Evaluation of 3 methods (ASigNTF, ABIS, CIBERSORT) and 4 signatures (Signature 415, Signature 915, ABIS, CIBERSORT) on a subset from GSE118165 dataset corresponding to Immature, Mature and Memory NK cells purified by FACS. Cell proportions (X-axis) for three cell types with highest estimated proportions are presented for each signature.

Since theoretically each sample represents a pure cell type obtained by flow cytometry, the correlation and *concordance correlation coefficients* [23] traditionally used to evaluate the performance of deconvolution methods cannot be used for this dataset. To overcome this problem, the performance of each signature for each immune cell type was evaluated by looking at the 3 cell types with the highest estimates. Assuming that the correct cell type is among these 3 cell types, the signature receives a score of ‘low’, ‘medium’ or ‘high’, if the correct cell type corresponds to the lowest, medium or highest estimate. If the correct cell type is not among the first 3 cell types, it receives a score ‘absent’. For example, NK cells in immature NK samples have the highest proportion for signatures 415 and 915 and receive the highest score of ‘high’.

If the correct cell type is not part of the signature under evaluation, the signature’s score is considered as missing.

The scores are shown in Figure 11. Cell types (pDC, Gamma-delta VD2-, Gamma-delta VD2+, Plasmablasts) that could not be assigned to these five groups were considered separately as the group “Others”.

**Figure 11.**
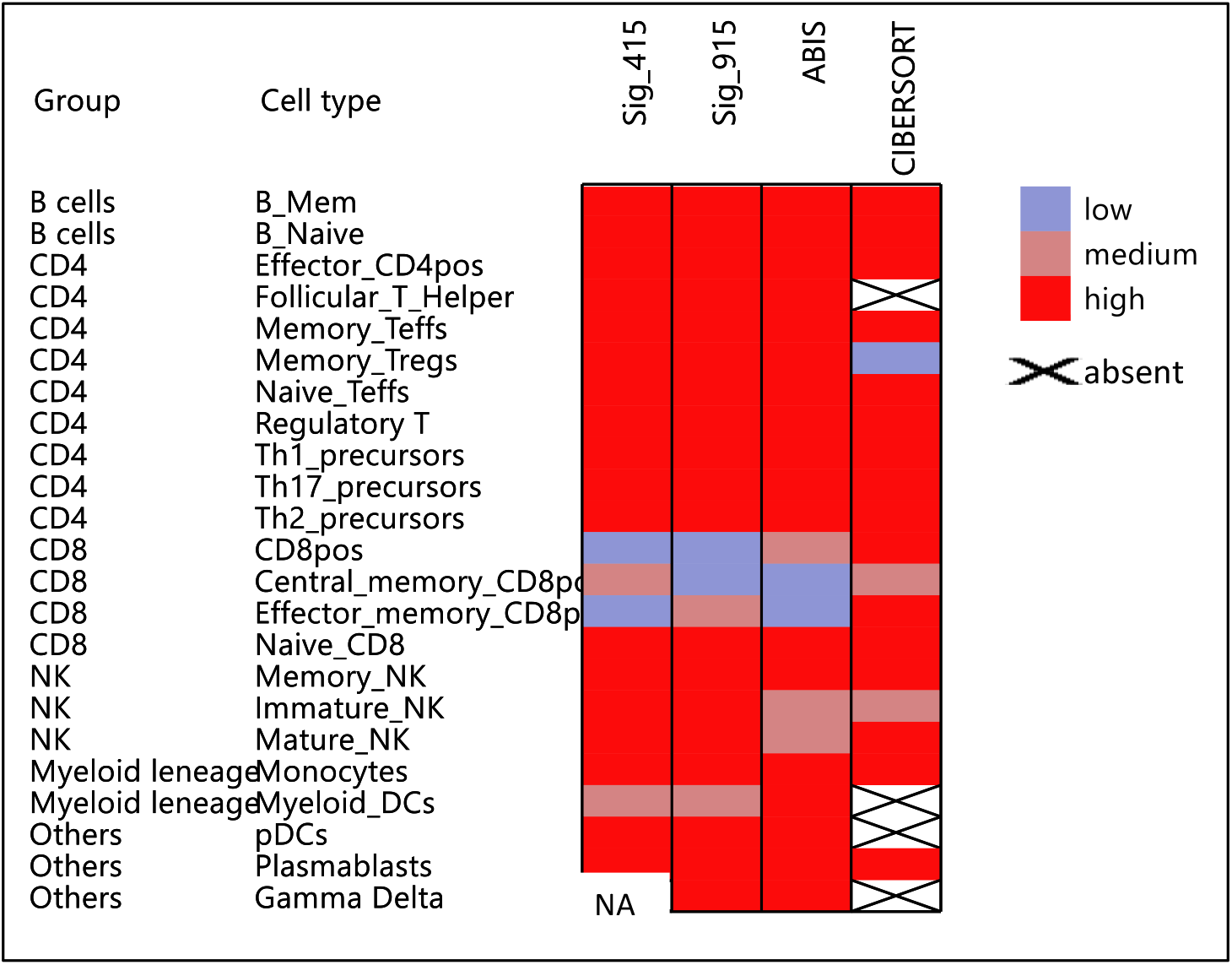
Heatmap summarizing the evaluation of 3 methods (ASigNTF, ABIS, CIBERSORT) and 4 signatures (Signature 415, Signature 915, ABIS, CIBERSORT) on whole RNA-seq dataset GSE118165 containing 166 samples corresponding to 25 immune cell types before or after stimulation. Each signature for each immune cell type is evaluated considering the 3 cell types with the highest estimated proportions. The signature receives a score of ‘low’, ‘medium’ or ‘high’, if the correct cell type corresponds to the lowest, medium or highest estimate. If the correct cell type is not among the first 3 cell types, it receives a score of ‘absent’.

To assess associations between scores, signatures and cell type groups, we performed a Multiple Correspondence Analysis, MCA [24]. ABIS-seq and our two signatures have similar performances on this dataset, surpassing CIBERSORT. In fact, ABIS-seq and our two signatures scored highest in four of the six cell groups. CIBERSORT was not able to correctly identify cell types from the myeloid lineage group and the group containing pDC and Gamma-delta cell types. In the MCA Plot (Figure 12), the ‘Other’ and Myeloid Parentage groups are associated with the CIBERSORT signature at the top of the Y-axis, as it has received an ‘absent’ score for most of the cell types belonging to these groups. In contrast, the 415, 915, and ABIS-seq signatures in these groups receive ‘medium’ to ‘high’ scores and are therefore away from the CIBERSORT signature in the lower part of the Y-axis, near the ‘high’ score. The CD8 group is associated with a ‘low’ score, since 10 of the 16 points in this group correspond to ‘low’ or ‘medium’. CIBERSORT scores slightly better on the cell types in this group compared to the 3 other signatures.

**Figure 12.**
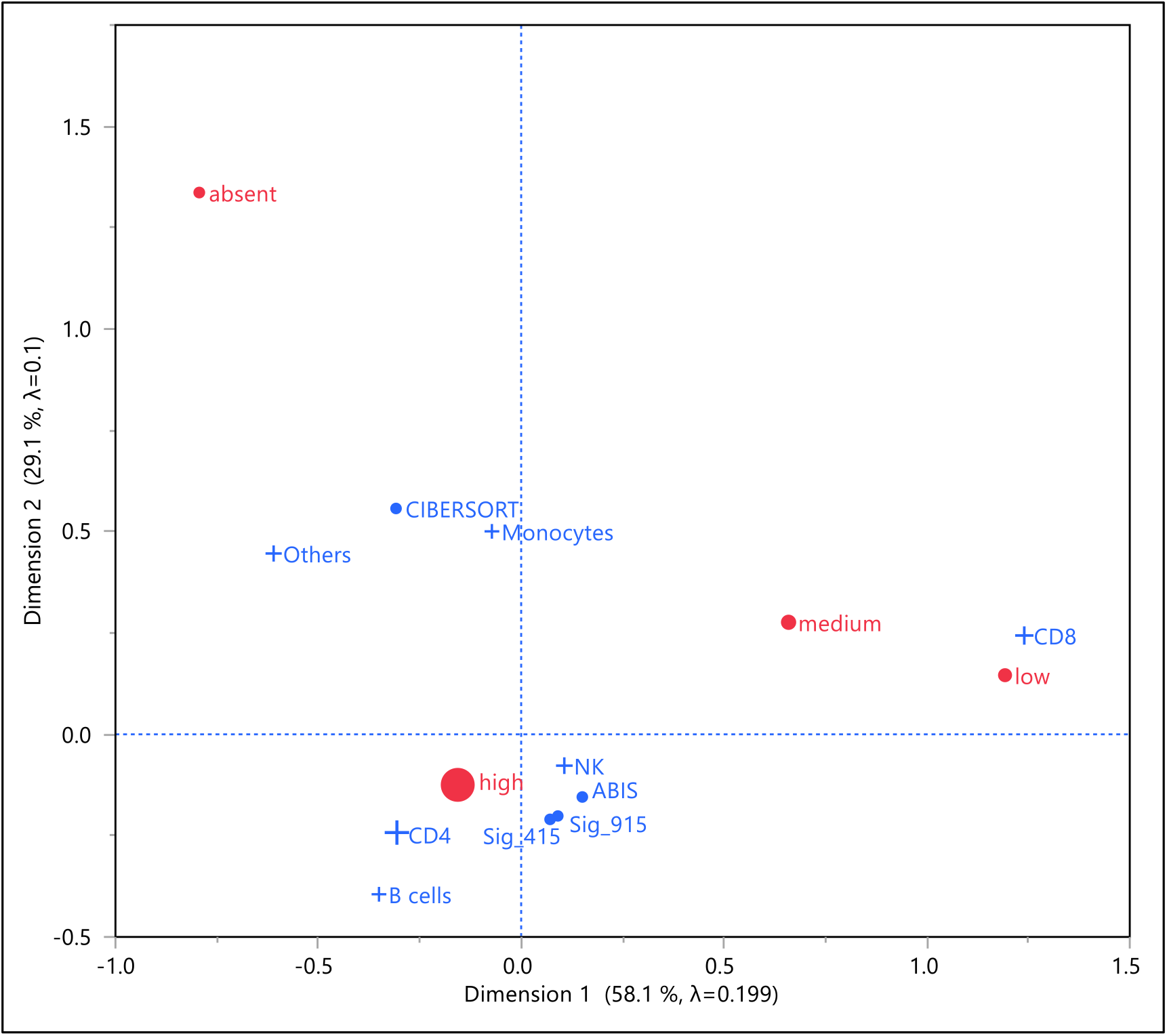
Multiple Correspondence Analysis of signature scores for RNA-seq dataset GSE118165. The size of each point corresponds to its frequency. ‘Low’, ‘medium’ and ‘high’ scores are distributed along the X-axis, while the Y-axis ranges between ‘absent’ and ‘high’. The CD8 group is associated with a ‘low’ score given that 10 out of the 16 scores for this group equal ‘low’ or ‘medium’. The groups B cells, CD4+ T cells and NK are associated with a ‘high’ score. The groups ‘Others’ and Monocytes are associated with the CIBERSORT signature in the upper part of the Y-axis, because of its ‘absent’ score for most of the cell types belonging to these groups. The signatures 415, 915 and ABIS-seq receive ‘medium’ to ‘high’ scores in these groups and are close to the ‘high’ score, in the lower part of the Y-axis. The signatures 915 or 415 outperformed other signatures for Immature, Mature and memory NK cells. CIBERSORT outperformed other signatures for the CD8 T cells group but lagged behind the CD4+ T cells and monocytes groups.

### Estimating immune cell types proportions from PBMC samples profiled by RNA-seq

To evaluate our signatures on different immune cells present in PBMC, we used the ABIS dataset (GSE107011), which contains transcriptomic signatures of PBMC from 13 Singaporean individuals (S13 cohort from [1]). In parallel, immune cells from these 13 PBMC samples were sorted by flow cytometry, which gave us access to the ground truth proportions of 29 cell types. Two measures were used: Pearson correlation coefficient (r) and Lin’s *concordance correlation coefficient* (ccc). The latter can be thought off as a correlation coefficient that enforces the correct scale and offset of the outputs [25]. The Pearson correlation coefficients and Lin’s concordance correlation coefficients are calculated per cell type.

Fourteen out of the 16 different immune subgroups were significantly correlated (p < 0.0001) between our signature 915 and flow cytometry (12/14 with our signature 415), including 10 subgroups with mean fractions below 5%. The box plots of correlations (r) and concordance correlation coefficients (ccc) show that the ABIS-seq signature performs better than the signature 415, closely followed by the signature 915, while the CIBERSORT signature lags far behind (Figure 13).

**Figure 13.**
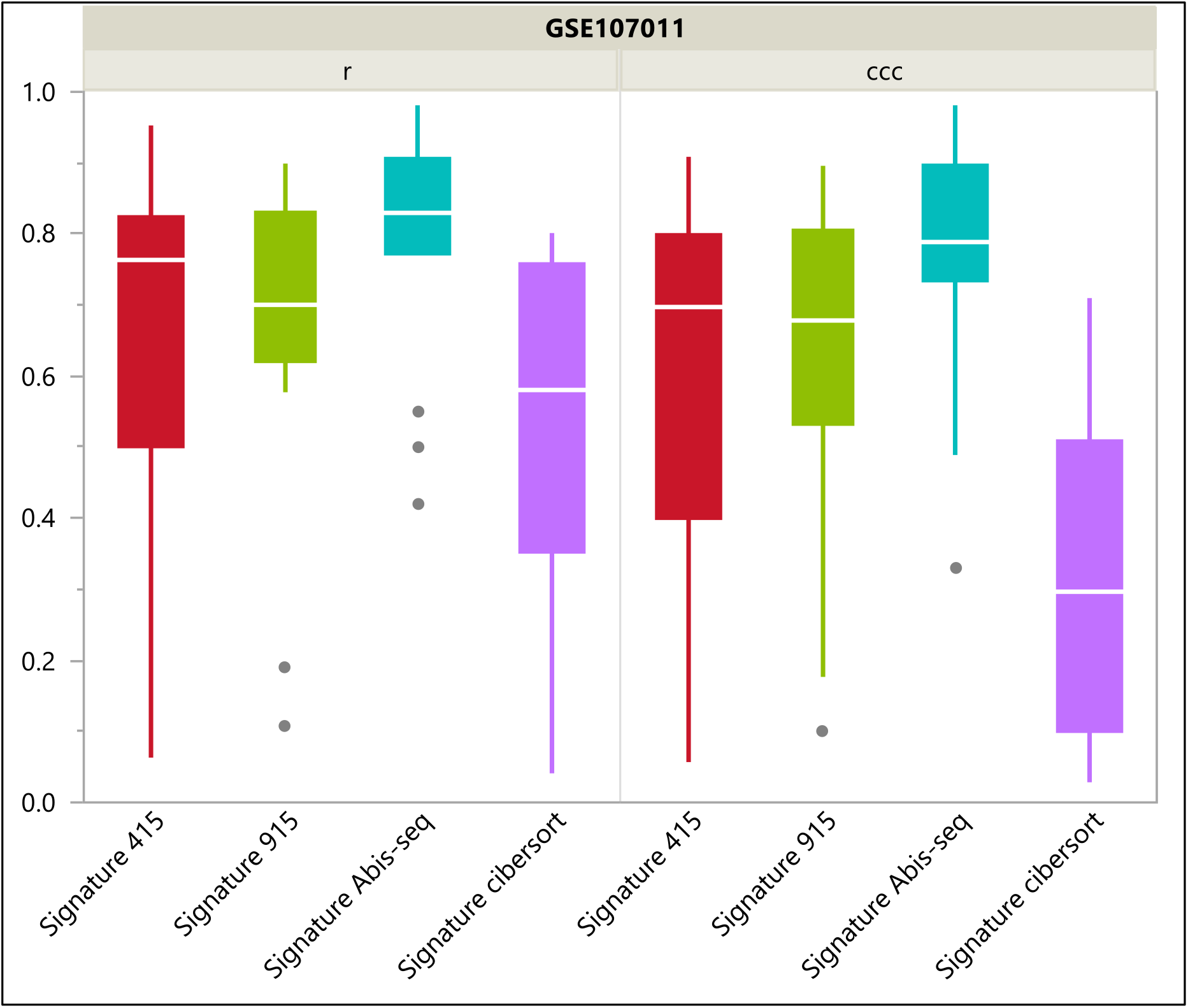
Box plots (study GSE107011) RNA-Seq dataset for 13 PBMC samples. Evaluation of 3 methods (ASigNTF, ABIS, CIBERSORT) and 4 signatures (Signature 415, Signature 915, ABIS, CIBERSORT) relative to flow cytometry proportions. Pearson correlation coefficients (r) and Lin’s concordance correlation coefficients (ccc) results are estimated per cell type and summarized by box plots for each signature.

The cell type grouping was chosen in the ABIS-seq signature to achieve the maximal concordance correlation coefficient (ccc) between the deconvolution estimates of the 13 PBMC samples and the proportions found by flow cytometry. It is therefore expected that this signature will outperform other signatures on this particular dataset. Noteworthy, we also used these samples to estimate abundance factors that are needed to correct nnls estimates for RNA abundance, which is cell type-specific, as described above. Since ccc values, unlike correlations, are based on corrected estimates, only the correlations should be regarded as the result of an external, unbiased validation, in the evaluation of our signatures.

### Estimating mixture proportions with unknown content

To compare the technical performance of our signatures with other methods for mixtures of unknown content, we used a benchmark dataset, GSE64385 [26], consisting of five admixed immune cell populations in combination with the colon cancer cell line HCT116. These 12 RNA mixtures were profiled by microarray and were therefore also of interest to assess whether our signatures could be used to deconvolute data from microarray technology. These mixtures with a colon cancer cell line simulate human solid tumors with different leukocyte infiltrations ranging from 1% to 48%.

We found that the fractions estimated by our signatures 415 and 915 for cell types present in mixtures were highly correlated with the known fractions and that the mean concordance correlation coefficients were similar for signature 415, signature 915 and ABIS-seq, followed by the CIBERSORT signature, although with a lower range for the latter (Figure 14).

**Figure 14.**
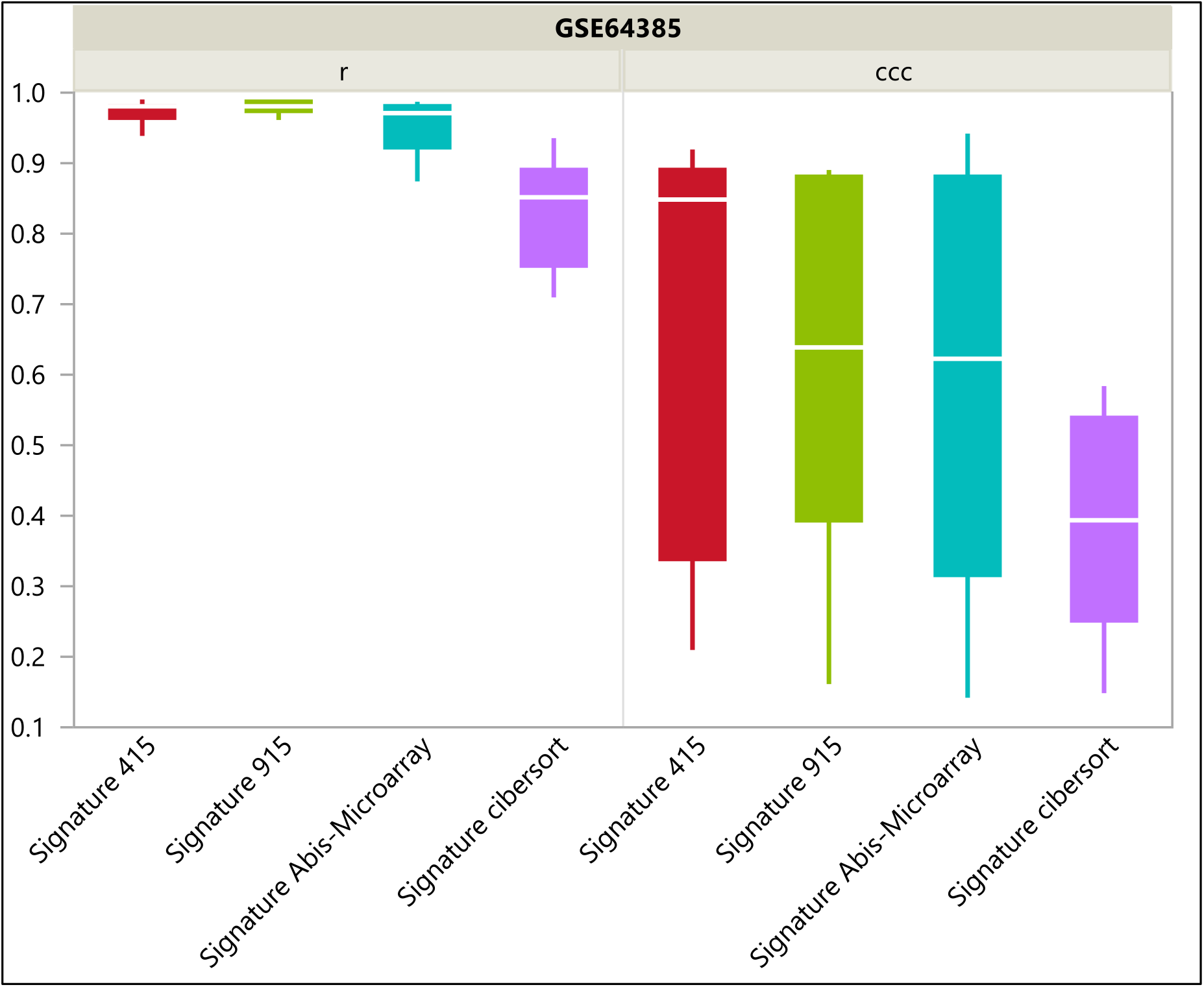
Box plots (study GSE64385) Microarray dataset for 12 mixtures including 6 cell types (B cells, T cells, NK cells, Monocytes, Neutrophils, colon cancer cell line HCT116). Evaluation of 3 methods (ASigNTF, ABIS, CIBERSORT) and 4 signatures (Signature 415, Signature 915, ABIS, CIBERSORT) relative to ground truth RNA content in ad-mixtures. Pearson correlation coefficients (r) and Lin’s concordance correlation coefficients (ccc) results are estimated per cell type and summarized by box plots for each signature.

Specifically, the mean correlation coefficient obtained for signatures 415 and 915 is about 0.98, ranging between 0.94 and 0.99, which implies the potential usefulness of our signatures for analyzing the composition of leukocytes from tumor samples profiled with microarray technology.

We also investigated a possible correlation between the real fraction level and the estimation error (Figure S2, suppl. material). With the exception of the B cells, no correlation with the estimation error was found for our signatures and the ABIS-seq signature, in contrast to the CIBERSORT signature, which showed a clear negative correlation in 4 of the 5 cell type groups studied – indicating that this signature might be less robust if the real fraction becomes too small.

### Estimating immune cell types proportions from PBMC samples profiled by microarray

Next, we used the GSE65133 dataset generated from a set of 20 PBMC samples obtained from adults of different ages who received influenza vaccination (NCT01827462). The gene expression profiling of these samples was performed by microarray and analyzed in parallel by flow cytometry to enumerate several leukocyte subsets [15].

Since this dataset and the signature matrices provide a different composition of cell types, we have validated only the 5 most important immune subgroups. Considering the Pearson correlation coefficient (r), the fractions estimated by our signature 415 in 4 different immune subgroups were significantly (p < 0.0001) correlated with flow cytometry. Interestingly, this study shows that our signature 415 enumerates cell type composition with a higher mean correlation (r) than the other three signatures, but with a mean concordance correlation coefficient (ccc) below ABIS-seq and CIBERSORT. This discrepancy is directly related to the factors used to correct for RNA abundance, which in this study, unlike other studies, resulted in lower ccc performance. This illustrates the difficulty of estimating correction factors that are valid in all experiments. In terms of the concordance correlation coefficient, ABIS-seq surpasses the other three signatures in mean, albeit not in range (Figure 15).

**Figure 15.**
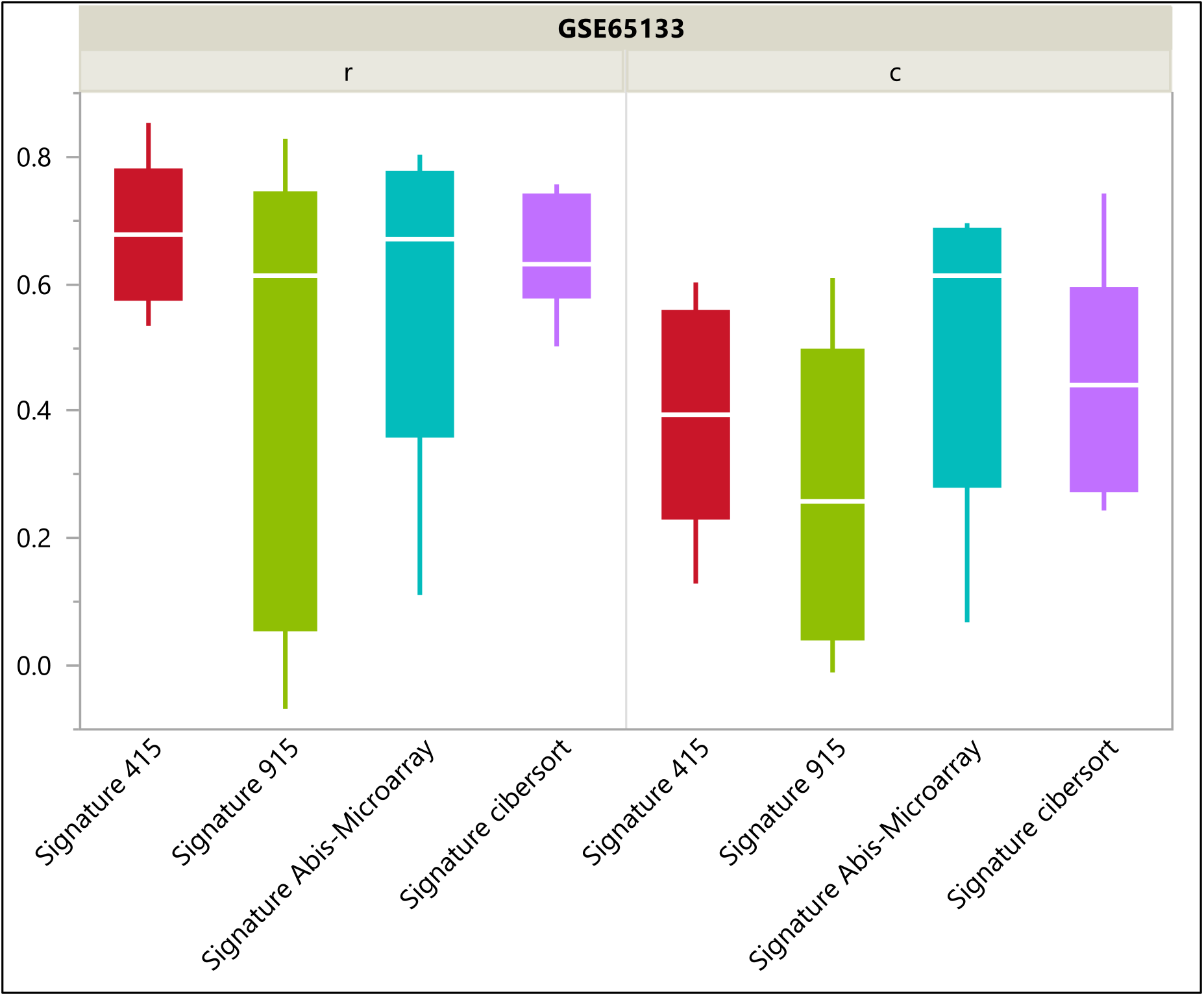
Box plots (study GSE65133) Microarray dataset with 20 PBMC samples. Evaluation of 3 methods (ASigNTF, ABIS, CIBERSORT) and 4 signatures (Signature 415, Signature 915, ABIS, CIBERSORT) relative to flow cytometry proportions. Pearson correlation coefficients (r) and Lin’s concordance correlation coefficients (ccc) are estimated per cell type and summarized by box plots for each signature.

## Discussion

The present study illustrates the usefulness of an agnostic, unsupervised approach to identify cell types or groups of cell types that are suitable for the deconvolution of complex datasets such as PBMC, whole blood or tumor samples infiltrated by immune cells. In fact, non-negative tensor factorization allows easy identification of cell types that have a detectable and specific transcriptomic fingerprint, helps to define cell types that are close to each other and which should be grouped for accurate deconvolution from transcriptomic data of bulk samples. We also propose two simple rules to guide the selection of marker genes. Together with our expression-based scaling scheme, our signature construction process results in well-conditioned matrices without the need to optimize their condition number. In contrast, using classical statistical tests does not guarantee the specificity of the marker genes, as we have shown in the present study. To improve specificity, marker genes must be filtered by minimizing the condition number of the signature matrix [1] [2] [6] [15] [21]. Nevertheless, the solution to this problem is NP-hard. A common approach is to rank genes by increasing significance and remove genes sequentially until the condition number is minimized [1]. However, since this significance-ranking does not fully reflect the specificity of the marker genes, this minimization scheme is not optimal. Beyond the benchmarked datasets, the much lower condition number of our signature 915 (36% lower than the CIBERSORT signature) indicates that it may be more reliable than other signatures. The scaling scheme used in our signatures ensures that all marker genes in the represented cell type are equally important, regardless of their expression level, thus avoiding additional filtering of genes with extremely high levels [1]. It is remarkable that this scaling scheme also plays an important role in the achieved low condition number of our matrices – which would otherwise be much higher. An interesting weighing scheme is used in [19], where genes that have less cross-cell type and more cross-subject variations have a low leverage effect, making pre-selection of marker genes unnecessary. However, the level of marker gene expression is not considered. We follow a more traditional approach by pre-selecting marker genes but recognize the need for less strict selection and therefore provide two alternative signatures with different amounts of marker genes.

The HHI was originally proposed by Herfindahl and Hirschman to assess the size of companies relative to the industry and provide an indicator of the degree of competition between them [16]. Newer versions of the HHI have been proposed, in particular the sparseness measure [27], which mathematically corresponds to the HHI, but lies between 0 (all entries in the vector are equal) and 1 (only one entry is not negligible). We decided to use the original HHI because of its good interpretability of reflecting the number of non-negligible entries in a vector.

We have observed in the GSE118165 study that the absolute values of cell type quantities returned by the ABIS package may be negative. If the aim of the study is to perform a cell type-based differential analysis, the estimates for each cell type must be normalized to ensure that the sum of the estimates is equal to 1 [28]. The presence of negative values returned by the ABIS package does not allow to perform the sum-to-1 normalization.

In summary, the application of ASigNTF is capable of identifying up to 16 major immune cell types suitable for deconvolution when applied on transcriptomics data from blood samples. We are indeed proposing two signatures, one of 415 genes for deconvolution of microarray data and the other of 915 genes for RNA-seq platform. Our signatures were benchmarked on external datasets with reference signatures (ABIS-seq, CIBERSORT). Both signatures outperformed CIBERSORT in the deconvolution of RNA-seq data. One possible reason is that CIBERSoRT is less robust for detection of rare cell populations, as it was demonstrated in the GSE64385 study. Our signature of 415 genes allows to detect 14 cell types compared to ABIS-microarray signature which is detecting only 11 cell types. This smaller signature may perform better than the larger signature of 915 genes on microarray data since, as it was determined in the GSE65133 study, almost 100% of its genes are detected by microarray probes, compared to about 50% of the genes from the signature of 915 genes. All in all, the choice of the signature for deconvolution can be influenced by several factors, such as the cell types present in the samples and the technology used for sample processing. Despite the good global performance of our signatures, some cell types may still be difficult to deconvolute. Therefore, we advocate the use of at least two different approaches for deconvolution due to their complementarity.

Although the deconvolution approach cannot replace the analysis of data at the single-cell level, it is more cost-effective because it allows the reuse of large transcriptomic data obtained by RNA-seq or microarrays and can be easily applied to large cohorts with thousands of samples. Computationally, deconvolution also requires fewer resources and can generate cell proportions for thousands of samples in several minutes, while the analysis of data at the single cell level is quite computationally intensive and time consuming.

In conclusion, we present in this article ASigNTF, a new universal and agnostic analysis strategy to estimate the frequency of different cell subtypes in transcriptomics data obtained from mixtures. We applied this strategy to data from blood samples (whole blood or PBMC) and demonstrated that the method works at least as well as reference methods from this field, such as CIBERSORT or ABIS. The results of this article are therefore manifold: 1) Our signatures are available and can be used to deconvolute data from whole blood or PBMC transcriptomics analyses (RNA-seq or microarray), 2) The proposed method is customizable as it can be refined with personal data (e.g. by adding further patients/samples) – NTF provides a model that can be easily updated in this respect. 3) The presented approach can be applied to other tissues – e.g. in oncology – and compared with MCP counters.

## Supporting information

Signature 415 genes

Signature 915 genes

## Availability

All the steps of our analysis: cell type grouping and construction of a signature, were implemented in an annotated Python notebook (suppl. material), which returns visualizations, e.g. heatmaps, PCA plots and box plots, and evaluates the marker gene criteria described in the paper. A separate notebook is provided to perform the deconvolution (suppl. material). To perform NTF, we used the package NMTF, which can be downloaded through the python pip install platform.

## Authors’ contributions

Galina Boldina and Paul Fogel equally contributed to the article.

- Galina Boldina participated in study conception, identified external datasets for benchmarking, performed the data interpretations and manuscript drafting.
- Paul Fogel designed the methods, performed the statistical analyses, wrote the manuscript draft.
- Corinne Rocher provided a critical review of the manuscript.
- Charles Bettembourg implemented Python notebooks using the methods presented in the article.
- George Luta revised the manuscript critically for important intellectual content.
- Franck Augé performed the bibliographic search and provided a critical review of the manuscript.

All authors read and approved the final manuscript.

## Conflicts of interests

None declared.

## Acknowledgements

We would like to thank Arnaud Droit for his careful review of our manuscript.

## Notes

### Competing Interest Statement

The authors have declared no competing interest.

