## Supplemental Figures for "ASigNTF: Agnostic Signature using NTF – a universal agnostic strategy to estimate cell-types abundance from transcriptomic datasets"

### Slide 1
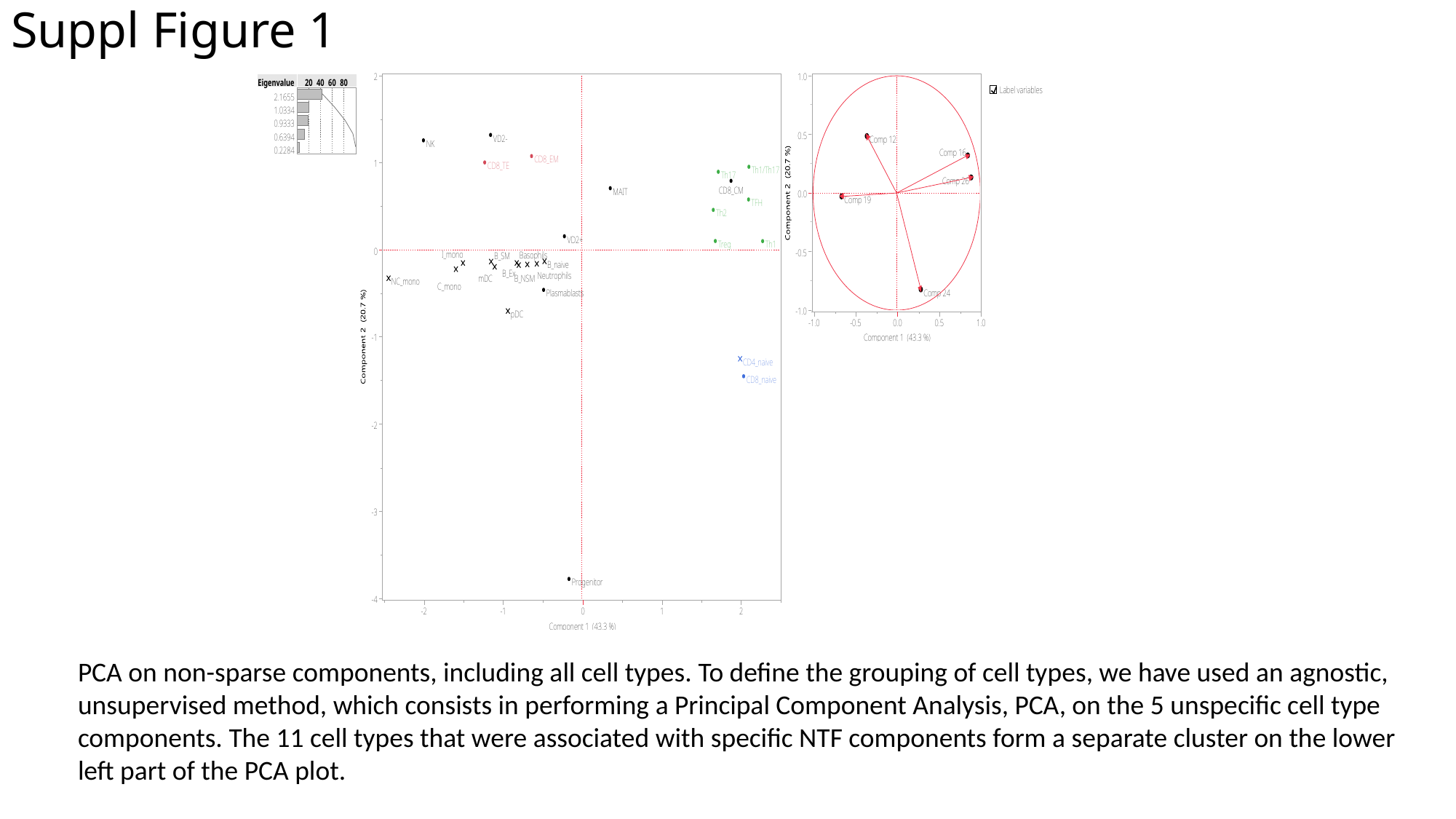

#### Suppl Figure 1
PCA on non-sparse components, including all cell types. To define the grouping of cell types, we have used an agnostic, unsupervised method, which consists in performing a Principal Component Analysis, PCA, on the 5 unspecific cell type components. The 11 cell types that were associated with specific NTF components form a separate cluster on the lower left part of the PCA plot.

### Slide 2
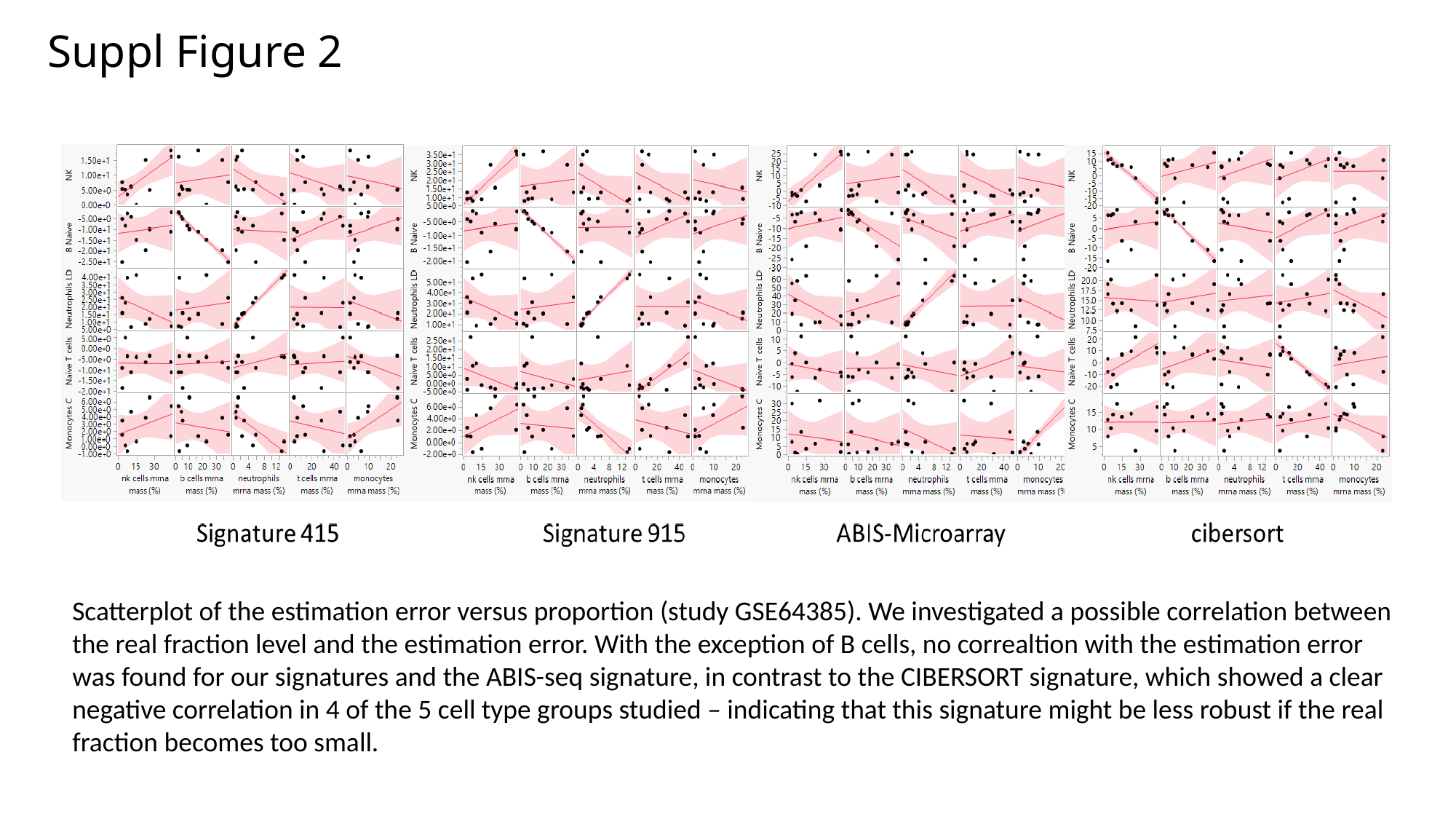

#### Suppl Figure 2
Scatterplot of the estimation error versus proportion (study GSE64385). We investigated a possible correlation between the real fraction level and the estimation error. With the exception of B cells, no correaltion with the estimation error was found for our signatures and the ABIS-seq signature, in contrast to the CIBERSORT signature, which showed a clear negative correlation in 4 of the 5 cell type groups studied – indicating that this signature might be less robust if the real fraction becomes too small.
